# SPIN90 modulates the architecture of lamellipodial actin in an ARPC5L dependent fashion

**DOI:** 10.64898/2025.12.01.691495

**Authors:** LuYan Cao, Angika Basant, Miroslav Mladenov, Naoko Kogata, Antoine Jegou, Guillaume Romet-Lemonne, Sophie Brasselet, Manos Mavrakis, Michael Way

## Abstract

The Arp2/3 complex in mammals, consists of 8 iso-complexes with different properties as there are two isoforms of the Arp3, ArpC1 and ArpC5 subunits. All iso-complexes still generate actin branches when activated by class 1 nucleation promotion factors. Using recombinant Arp2/3 iso-complexes with defined compositions, we now demonstrate that SPIN90 selectively activates ArpC5L- rather than ArpC5- containing complexes to generate linear actin filaments. Consistent with this, SPIN90 at the leading edge of migrating cells enhances the recruitment of ArpC5L-, but not ArpC5- containing, Arp2/3 complexes to lamellipodia. These SPIN90-Arp2/3-ArpC5L complexes generate linear actin filaments that integrate into lamellipodia and impact protrusion efficiency. Moreover, using polarized light microscopy, we show that loss of SPIN90 leads to changes in the lamellipodial actin filament architecture. Our results demonstrate that SPIN90 regulates the organisation and dynamics of lamellipodial actin in an Arp2/3 iso-complex dependent fashion.

**Significance statement:** Recent studies have shown that mammalian Arp2/3 complexes exist as eight iso-complexes with different molecular and cellular functions as three of the seven subunits are encoded by two different genes. However, all known NPF-I activators, including N-WASP and WAVE, activate these Arp2/3 iso-complexes albeit with different efficiencies to promote branched actin assembly. Here, we report SPIN90 is the first Arp2/3 activator that selectively promotes the recruitment and activation of ArpC5L- but not ArpC5- containing Arp2/3 complexes. These findings provide new insights into how distinct Arp2/3 complexes may be differentially controlled in cells.

## Introduction

Highly conserved throughout evolution, the Arp2/3 complex is a unique actin nucleator, essential for generating branched actin networks (1–4). Consequently, the Arp2/3 complex is an important regulator of the architecture and dynamics of the actin cytoskeleton (5–9). Activated by class I nucleation-promoting factors (NPFs), such as WAVE, N-WASP, WASP, and WASH, the Arp2/3 complex can bind to pre-existing actin filaments, also known as mother filaments, and initiate the formation of a new actin branch, or a daughter filament (9–11). Alternatively, when activated by SPIN90 (also known as WISH, NCKIPSD, or Dip1 in yeast), the Arp2/3 complex is unable to bind to an existing actin filament and rather generates a new linear actin filament (8, 12). Given its ability to nucleate actin networks, the Arp2/3 complex is indispensable for various cellular functions, including endocytosis, cell migration, and DNA repair (2, 13–17). Among these processes, SPIN90–Arp2/3 contributes to the initiation of yeast endocytosis by providing the actin filaments required for actin branch formation (17) and in the regulation of mammalian cortical actin by tuning its architecture (13, 18). It has also recently been reported that SPIN90-Arp2/3 plays a role in focal adhesion maturation (19).

The Arp2/3 complex contains seven subunits, the <u>A</u>ctin <u>R</u>elated <u>P</u>roteins Arp2 and Arp3, as well as ArpC1-ArpC5 (1, 3, 20). In mammals, there are two isoforms of Arp3, ArpC1, and ArpC5, encoded by two different genes (1, 21, 22). In humans, Arp3 and Arp3B are 91% identical, while ArpC1A/ArpC1B and ArpC5/ArpC5L each share 67% identity (23). These different subunit isoforms have been shown to give the Arp2/3 complex distinct cellular and physiological functions (11, 24–27). In addition, mutations in human ArpC1B cause severe inflammation and immunodeficiency (28, 29). Loss of ArpC5 also leads to multiple congenital deficiencies, inflammation, dysregulated host-microbiome interactions, and increased mortality (30–32).

To understand the molecular mechanism underlying these phenotypes, it is essential to dissect the properties of the different Arp2/3 iso-complexes. Arp2/3 iso-complexes have subtly but recognizably different structural conformations (26, 33, 34) and respond differently to the widely used Arp2/3 inhibitors, CK-666 and CK-869 (35). Previous studies have also shown that all Arp2/3 iso-complexes, regardless of their subunit composition, can generate actin branches. Complexes containing ArpC1B or ArpC5L are, however, more efficient at branch formation than those with ArpC1A or ArpC5 (23), while networks generated by Arp3B-containing complexes disassemble faster than those formed by Arp3-containing complexes in cells due to oxidation of Methionine 293 of Arp3B by MICAL2 (36).

In contrast, it is still unclear whether all Arp2/3 iso-complexes generate linear actin filaments when activated by SPIN90. Moreover, our previous work demonstrated that although CK-666 inhibits all Arp2/3 iso-complexes from generating linear filaments, it only blocks ArpC1A containing complexes producing actin branches (35). These findings suggest that different Arp2/3 iso-complexes engage fundamentally distinct mechanisms depending on whether activation drives linear filament formation or actin branching. Our structural analysis using recombinant Arp2/3 showed the SPIN90 can generate linear actin filaments with ArpC1B and ArpC5L containing complexes (37). However, the structure of the SPIN90–Arp2/3 complex, which contains Arp2/3 complex purified from bovine thymus, does not have sufficient side-chain resolution to determine isoform-specific information (12, 38). Hence, it is not known if ArpC1A and/or ArpC5 containing complexes can also generate linear filaments.

In this study, using in vitro biochemical approaches, we found that only ArpC5L- containing Arp2/3 complex can generate linear actin filaments when activated by SPIN90. In cells, SPIN90 accumulates at the leading edge of lamellipodia, where it recruits and activates ArpC5L-, but not ArpC5-containing Arp2/3 complexes. These linear filaments nucleated by SPIN90-Arp2/3-ArpC5L complexes shape the architecture of lamellipodial actin, impacting actin dynamics and leading-edge protrusion.

## Results

### SPIN90 preferentially activates ArpC5L-containing Arp2/3 complexes

Using purified recombinant human Arp2/3 iso-complexes with defined compositions, we performed pyrene actin polymerisation assays to examine their capacity to accelerate actin polymerisation when activated by SPIN90. We found that, in the presence of SPIN90, Arp2/3 complexes containing ArpC1A-ArpC5L or ArpC1B-ArpC5L significantly increase the rate of actin polymerization compared to controls lacking SPIN90 (Fig. 1A). In contrast, Arp2/3 complexes containing ArpC1A-ArpC5 or ArpC1B-ArpC5 only induced actin polymerisation at levels slightly above controls when activated by SPIN90. These differences suggest SPIN90 only efficiently activates Arp2/3 complexes containing ArpC5L to generate linear filaments.

**Figure 1.**
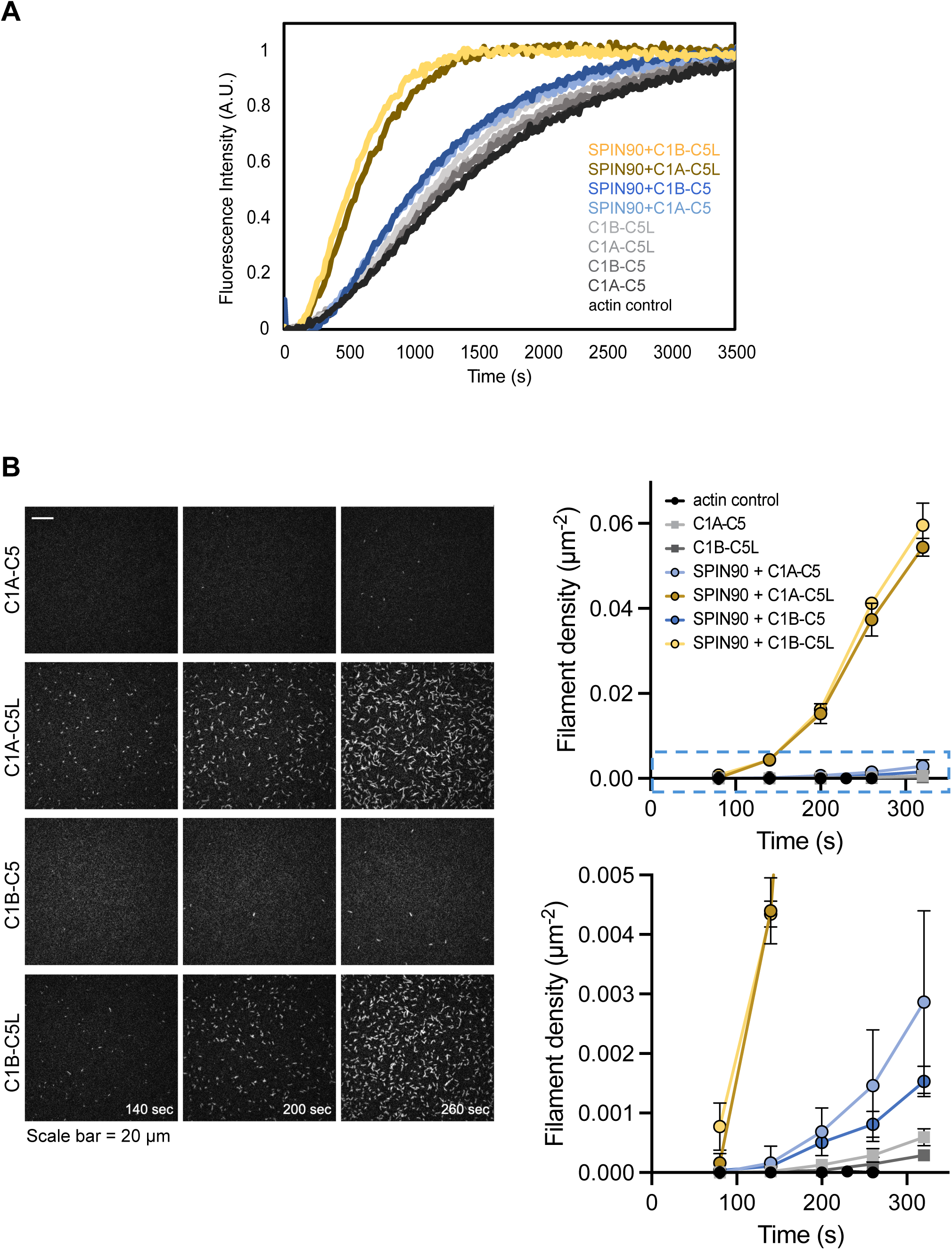
SPIN90 selectively activates ArpC5L containing Arp2/3 complexes. (A) SPIN90 efficiently activates ArpC5L but not ArpC5 containing Arp2/3 complexes. Representative plots of pyrene-actin polymerization assays with 10 nM Arp2/3 complex, 100 nM SPIN90 and 2.5 µM actin (5 % pyrene labelled). It demonstrates that ArpC5L- but not ArpC5-containing Arp2/3 complexes activated by SPIN90 induce rapid actin polymerization, regardless of the ArpC1 isoform. The experiments were repeated independently three times and yielded similar results. (B) Representative TIRF images demonstrate that ArpC5L- but not ArpC5-containing complex activated by SPIN90 efficiently generate actin filaments. A mixture of 20 nM Arp2/3 complex, 200 nM SPIN90 and 0.5 µM actin (15 % Alexa 488 labelled) was observed overtime. The upper graph shows quantitative analysis of filament density over time from three independent technical replicates. The lower graph shows the information in the zoomed-in box highlighted in the upper graph. The points indicate the mean density, and the error bars represent the standard deviation.

To confirm this, we performed TIRF microscopy to directly observe the generation of linear actin filaments by the different Arp2/3 iso-complexes in the presence of SPIN90. Actin alone, or in combination with Arp2/3 complexes, does not efficiently nucleate filament formation (Fig. 1B). When incubated with SPIN90, Arp2/3 complexes containing ArpC5, whether paired with ArpC1A or ArpC1B, had an extremely limited increase in actin nucleation. In contrast, Arp2/3 complexes containing ArpC5L, regardless of the ArpC1 isoform, produce linear filaments with 20- to 40-fold greater efficiency in the presence of SPIN90 (Fig. 1B). These results clearly show that linear actin filaments are generated by SPIN90-activated ArpC5L-containing Arp2/3 complexes.

### SPIN90 has a higher affinity for Arp2/3 complexes containing ArpC5L

Our finding that only ArpC5L-containing complexes can be efficiently activated by SPIN90 raises the question: what is the molecular basis for the differing efficiencies in linear filament nucleation among Arp2/3 iso-complexes? The recent structure of SPIN90 in complex with Arp2/3 reveals it binds directly to Arp2, Arp3, ArpC2 and ArpC4, as well as to the ArpC5/ArpC5L subunit, which have 67% identity (Fig. S1A, B) (12, 37, 38). Nevertheless, the resolution of current available structure is insufficient to resolve the side chain and N-terminus of ArpC5 subunit (12, 38). Consequently, it remains unclear whether ArpC5 or ArpC5L is present in the bovine Arp2/3 complex interacting with SPIN90 (Fig. S1C). In contrast and consistent with our biochemical observations, in the structure of SPIN90-with defined recombinant human Arp2/3 complex, it interacts with ArpC5L (37).

Given our observations, we wondered whether the ArpC5 subunit incorporation determines the affinity between SPIN90 and the Arp2/3 complex. We tried several approaches to examine the affinity between Arp2/3 iso-complexes and SPIN90. However, surface plasmon resonance, microscale thermophoresis and isothermal titration calorimetry failed to yield reproducible results. We therefore examined the binding between Arp2/3 iso-complexes and SPIN90 using GST pull-down assays. We found that GST-SPIN90 pulled down more Arp2/3 complexes containing ArpC5L compared to those with ArpC5, regardless of the ArpC1 subunit isoform (Fig. S1D).

To quantify the affinity of SPIN90 for Arp2/3 complexes containing either ArpC5L or ArpC5, we performed GST pull-down assays (39, 40) on Arp2/3 complexes containing ArpC1B/ArpC5 or ArpC1B/ArpC5L (Fig. 2A, B). Analysis of the unbound fraction of Arp2/3 following the incubation of a constant amount of the complex with increasing amounts of GST-SPIN90 bound to beads revealed that SPIN90 binds the ArpC1B/ArpC5L- containing Arp2/3 complex with a dissociation constant (K_D_) of 6.3 µM (95% CI, 5.2-7.5) (Fig. 2A). In contrast, the low amount of ArpC5-containing complex pulled down with GST-SPIN90 prevented reliable fitting so a dissociation constant (K_D_) could not be determined. Alternatively, incubation of a fixed amount of GST-SPIN90 beads with varying concentrations of Arp2/3 yielded a similar dissociation constant of 6.1 µM (95% CI, 2.6-20.1) for the ArpC1B/ArpC5L-containing Arp2/3 complex and SPIN90 (Fig. S1E). As before the affinity of the ArpC1B/ArpC5-containing Arp2/3 complex for SPIN90 remained unmeasurable. Further pulldown assays, however, revealed that ArpC5 (K_off_ = 0.067 min^-1^) and ArpC5L (K_off_ = 0.045 min^-1^) containing complexes exhibit similar dissociation rate from SPIN90 (Fig. 2B).

**Figure 2.**
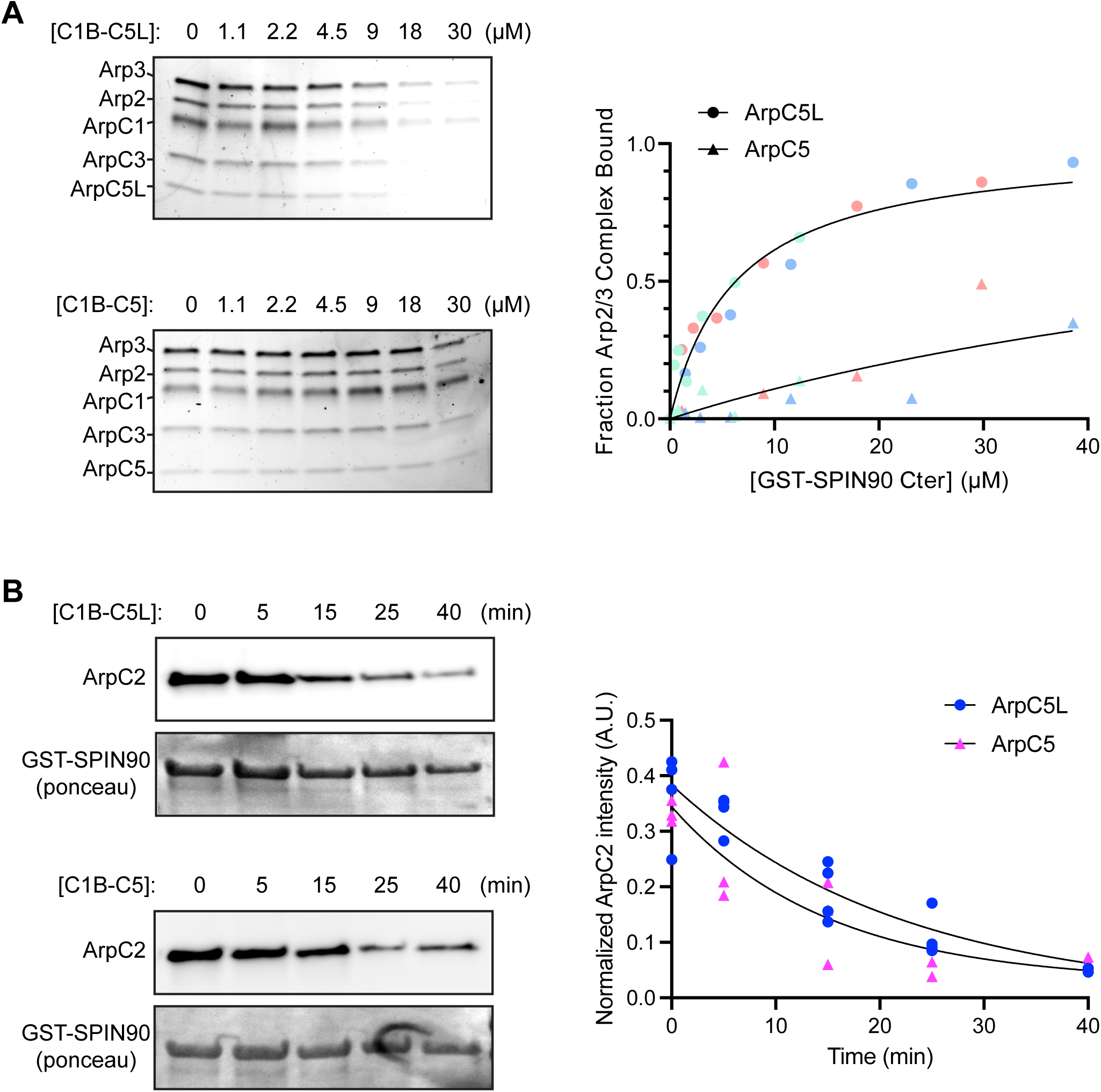
SPIN90 has a higher affinity for Arp2/3 complexes with ArpC5L than ArpC5. (A) Stain-free SDS-PAGE shows the amount of Arp2/3 complex remaining in the supernatant after incubating the same amount of Arp2/3 complex with increasing concentrations of SPIN90-attached beads. The graph shows the quantification of Arp2/3 fraction bound to SPIN90 versus the concentration of SPIN90 from three independent replicates, shown in different colour. The data was fit with one site-specific binding equation. (B) Immunoblots using an ArpC2 antibody show the amount of Arp2/3 complex attached to SPIN90 different time points after free Arp2/3 complexes were washed out. The graph shows the quantification of average normalised ArpC2 from three independent replicates. The data was fit with single exponential equation.

These results confirmed that the ArpC5 subunit isoforms have a strong impact on the association of SPIN90 for Arp2/3 iso-complexes. Furthermore, the distinct affinity may partially explain why SPIN90 effectively activates only ArpC5L-containing Arp2/3 complexes to generate linear actin filaments.

### SPIN90 is enriched at the leading edge

Another key question is the physiological impact of the selective interaction between SPIN90 and ArpC5L containing Arp2/3 complexes in cells. To study the impact of SPIN90 and Arp2/3 on the cellular actin, we used B16 cells which have large and constant lamellipodial branched actin networks (41, 42). We began by assessing the expression level of ArpC5 and ArpC5L containing Arp2/3 complexes in B16 cells using quantitative immunoblot analysis of recombinant mouse ArpC5 and ArpC5L subunits as standards (Fig. 3A). We found that ArpC5 and ArpC5L are both expressed in B16 cells, with ArpC5L levels being ∼1.7-fold higher than ArpC5.

**Figure 3.**
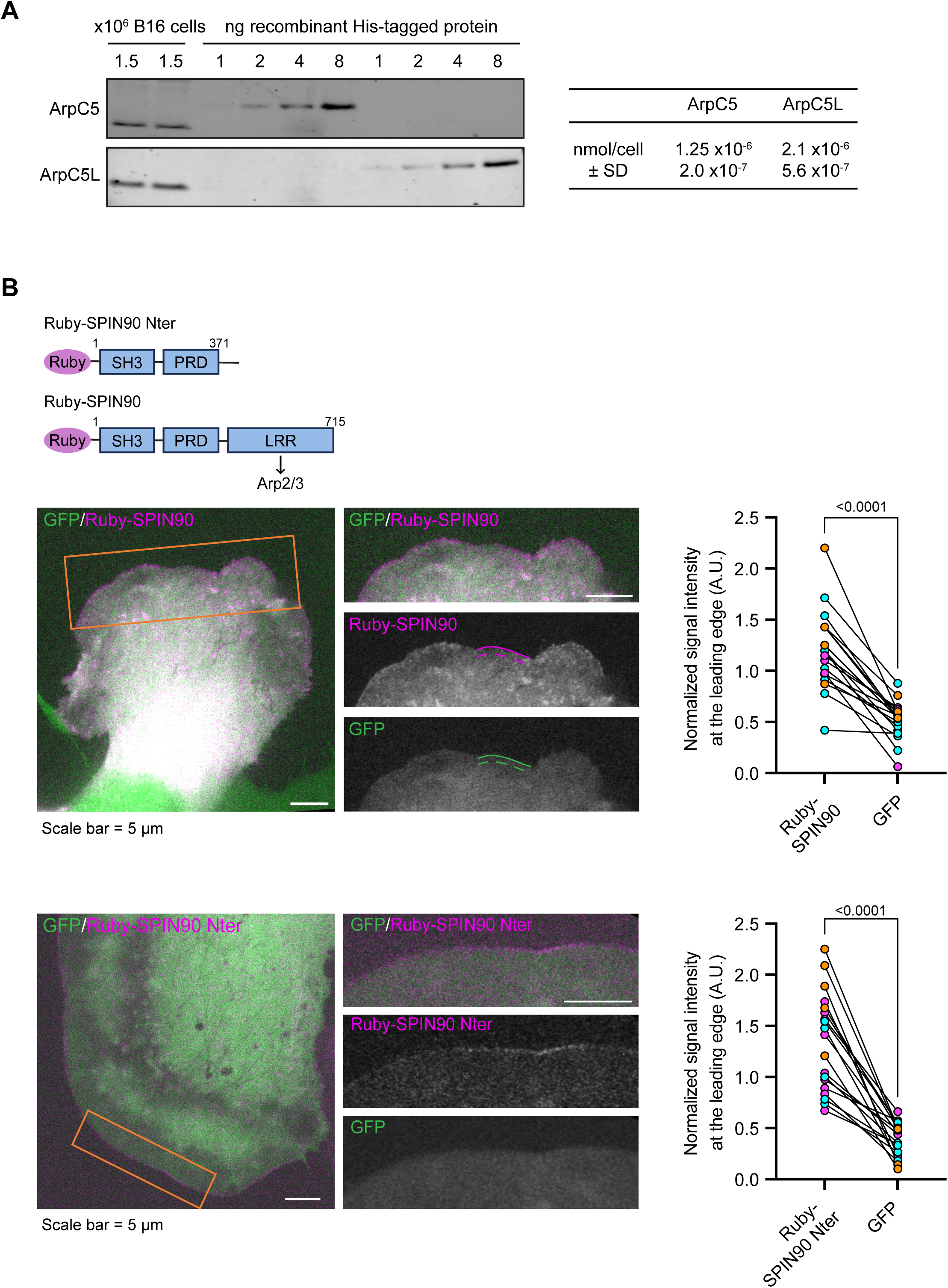
SPIN90 localizes at the leading edge and integrates into lamellipodial actin networks. (A) Quantitative immunoblot analysis of endogenous levels of ArpC5 isoforms in B16 cells using recombinant His-tagged ArpC5 and His-tagged ArpC5L as standards. The average concentrations and standard deviations from three independent replicates are shown in the table on the right. (B) Schematic shows the two Ruby-tagged SPIN90 constructs expressed in cells. Representative live images show B16 cells overexpressing GFP (volume marker) together with Ruby–SPIN90 (upper) or with Ruby-SPIN90 Nter (lower). Signal intensity (right) is quantified by normalizing the intensity at the leading edge (solid line) compared to that 2 µm from the leading edge (dashed line). Each pair of points represents measurements from the same cell with the different colours representing data from three independent replicates. A two-tailed paired t-test is used to estimate the p-value.

To determine the localisation of SPIN90 in B16 cells, we performed immunofluorescence analysis with both commercial and homemade SPIN90 antibodies but unfortunately none worked. We therefore expressed Ruby-tagged SPIN90 to visualize the localization of SPIN90 in live cells. We found that Ruby-SPIN90 is predominantly cytoplasmic, but the protein is enriched at the leading edge of lamellipodia which is not observed with the GFP internal control (Fig. 3B, Movie S1). A similar accumulation at the leading edge was observed for a Ruby-SPIN90 Nter construct that lacks the Arp2/3 interacting C-terminal Leucine Rich Region (LRR) of SPIN90 (Fig. 3B, Movie S2). This localisation demonstrates that the N-terminus of SPIN90 responds to an upstream signal responsible for determining the localisation of SPIN90.

### Both SPIN90 and Arp2/3 integrate into lamellipodial actin

To visualize ArpC5L and SPIN90 at the same time we expressed Ruby-SPIN90 in B16 cell lines in which NeonGreen is linked to the N-terminus of ArpC5L at its endogenous gene locus (Fig. S2A). We cannot distinguish ArpC5L bound to SPIN90 from ArpC5L located at actin branches, but we observed that both ArpC5L and SPIN90 accumulate at the leading edge of lamellipodia (Fig. 4A, B, Movie S3). Interestingly, during live imaging, we observed that both proteins initially appear at the leading edge and then move toward the cell body before disappearing (Movie S3). Hence, the trajectories of NeonGreen-ArpC5L and Ruby-SPIN90 display similar patterns, characterized by straight lines perpendicular to the leading edge (Fig. 4A). This is the first time such retrograde flow of SPIN90 has been seen in lamellipodia, although the rearward flow of Arp2/3 complexes has been described (41).

**Figure 4.**
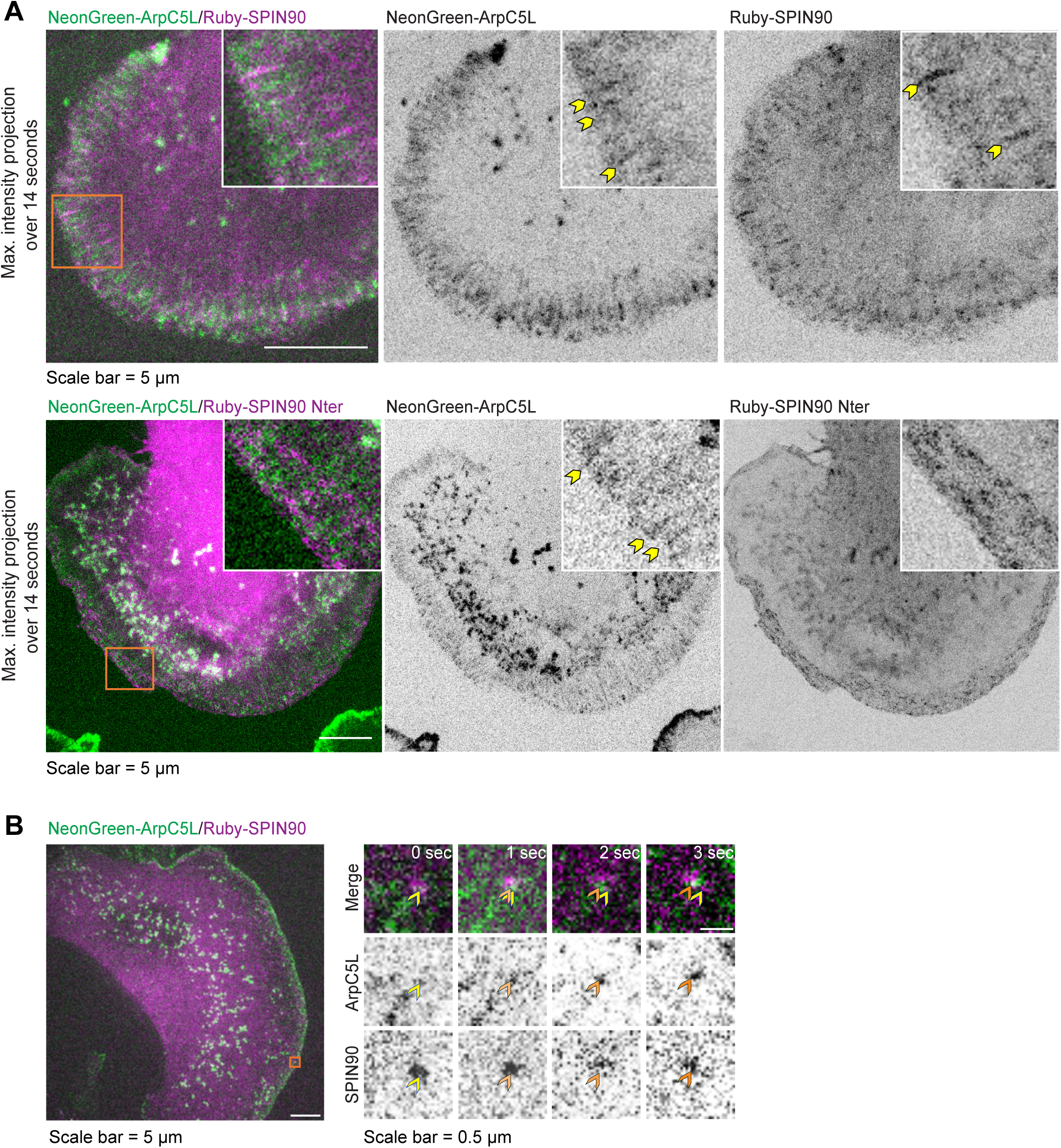
SPIN90 integrates into the lamellipodia with Arp2/3 complex. (A) Representative maximum intensity projection over 14 seconds for cells expressing endogenous NeonGreen–ArpC5L, together with overexpressed Ruby– SPIN90 or Ruby–SPIN90-Nter. ArpC5L and Ruby–SPIN90, but not Ruby–SPIN90-Nter undergo rearward movement in the lamellipodium (highlighted with yellow arrows). The similar trajectory pattern observed in all the cells (20 for SPIN90-FL and 13 for SPIN90-Nter) across three independent replicates. (B) A representative live image of B16 cells with endogenously labelled NeonGreen-ArpC5L and overexpressing Ruby–SPIN90. Right panel shows a zoomed-in image of the ROI highlighted in the orange box in the left panel. Yellow arrows indicate the exact position where ArpC5L and SPIN90 signals overlap at time 0, while orange arrows show the ArpC5L and SPIN90 signals moving together directionally over time. More examples are show in Fig. S2.

Given that most ArpC5L is at actin branch points and the existence of unlabelled ArpC5L, the SPIN90 and ArpC5L do not always overlap (Fig. A4). Despite that, we see examples in which that SPIN90 and ArpC5L colocalise at the leading edge and move towards the cell body together over a few seconds (Fig. 4B, S2, Movie S3). To determine whether SPIN90 integrates into the actin network through its interaction with Arp2/3 complexes, we examined the behaviour of Ruby-SPIN90 Nter in B16 cells expressing endogenous NeonGreen-ArpC5L. Live cell imaging reveals that while NeonGreen-ArpC5L undergoes retrograde flow, Ruby-SPIN90 Nter remained at the leading edge and did not move inward toward the cell body (Movie S4). The trajectories of NeonGreen-ArpC5L and Ruby-SPIN90 Nter exhibit distinct patterns (Fig. 4A). Taken together, our observations indicate that by binding to Arp2/3 complexes, SPIN90 integrates into lamellipodial networks and undergoes retrograde flow.

### SPIN90 enhances recruitment of ArpC5L but not ArpC5 to the leading edge

Our in vitro data demonstrate that SPIN90 interacts much better with Arp2/3 complexes containing ArpC5L, than those with ArpC5. To examine if SPIN90 enhances the recruitment of ArpC5L-containing Arp2/3 complexes, we quantified the relative level of NeonGreen-ArpC5L at the leading edge in the presence and absence of SPIN90 (Fig. 5A, S3A). We observed that in SPIN90 depleted cells, the relative level of NeonGreen-ArpC5L signal at the leading edge is reduced compared to control cells. When ArpC5 is knocked down, we observed increased recruitment of NeonGreen-ArpC5L to the leading edge, to compensate for the absence of ArpC5-containing complexes. Importantly, the depletion of SPIN90 in the absence of ArpC5 still results in reduced recruitment of ArpC5L to the leading edge (Fig. 5A). In contrast to ArpC5L, the relative signal of endogenous NeonGreen tagged ArpC5 at the leading edge does not negatively correlate with the depletion of SPIN90 as it slightly increases (Fig. 5B). The depletion of ArpC5L has little impact on the level of ArpC5 at the leading edge. However, the double knockout of ArpC5L and SPIN90 leads to its increase at the leading edge. Our observations on protein localisations were not due to changes in protein levels as immunoblots demonstrated that SPIN90 depletion does not affect the level of ArpC5 isoforms, while depletion of ArpC5 or ArpC5L had little impact on the level of the other isoform (Fig. S3A, B).

**Figure 5.**
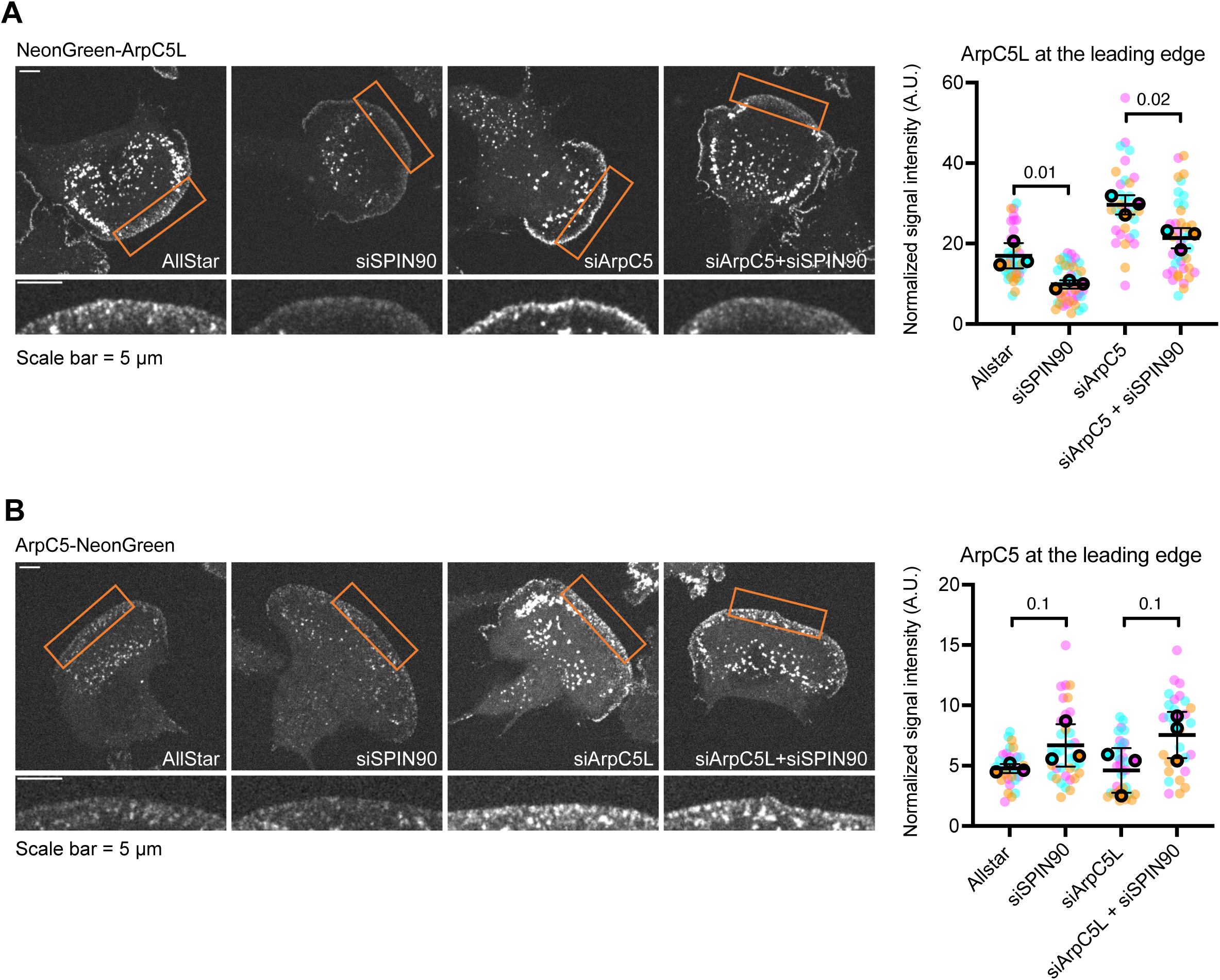
SPIN90 enhances the recruitment of ArpC5L but not ArpC5 to the leading edge. Representative images of live cells expressing endogenous NeonGreen–ArpC5L (A) or ArpC5-NeonGreen (B) after treatment with the indicated siRNAs. Allstar represents siRNA control. The graphs show quantification of the ArpC5L or ArpC5 signal intensity at the leading edge normalised with the expression level of NeonGreen tagged protein in the different conditions. Each point represents an individual cell, with different colours indicating 3 independent experimental replicates. For each repeat, 8-15 cells were imaged and quantified. The mean of each experiment is shown as a circle, and error bars are standard deviation. A two-tailed unpaired t-test was used to calculate the p-value.

To rule out the possibility that our observations were due to the side effect of the NeonGreen tag on the ArpC5/ArpC5L subunit, we performed immunofluorescence on siRNA-treated B16 cells. As expected, depletion of ArpC5 led to a decrease in the ArpC5/ArpC2 ratio. We observed that the ArpC5/ArpC2 ratio increased in both ArpC5L- and SPIN90-depleted cells (Fig. S3C). These observations support the hypothesis that SPIN90 enhances the recruitment of ArpC5L, but not ArpC5 containing Arp2/3 complexes to the leading edge.

### SPIN90 regulates lamellipodial actin architecture via ArpC5L

Arp2/3-mediated branches are geometrically constrained to form at an angle of approximately 70 degrees (7). However, it has also been reported that Arp2/3 complexes containing either ArpC5 or ArpC5L have a distinct impact on the architecture of lamellipodial actin networks although the mechanistic basis for this difference has not been established (26). Given that SPIN90 generates linear filaments only with ArpC5L- containing complexes, we wondered what impact its presence has on the architecture of lamellipodial actin networks, which may clarify the mechanistic basis for this difference By exploiting the sensitivity of polarized light excitation to the orientation of fluorophores attached to the actin filaments, polarized fluorescence microscopy can be used to assess actin filament organization (43, 44). It also allows the acquisition and processing of large amounts of cellular data quickly compared to cryo-electron microscopy methods. Using this approach, we quantified the organization of actin filaments in lamellipodia, excluding microspikes, in Alexa-488 phalloidin-stained cells. The Alexa-488 conjugate in phalloidin is constrained in mobility and is parallel to the axis of the actin filament helix (43). Thus, the orientation of Alexa-488 is a direct readout of actin filament mean orientation (angle rho, ρ) within a scale of ∼200nm (i.e. the optical resolution limit, Fig. 6A, B). Its angular spread around the mean orientation at each pixel is a complementary readout of actin filament alignment (angle psi, ψ) at the molecular level, within the scale of ∼200 nm (Fig. 6A, S4A-C).

**Figure 6.**
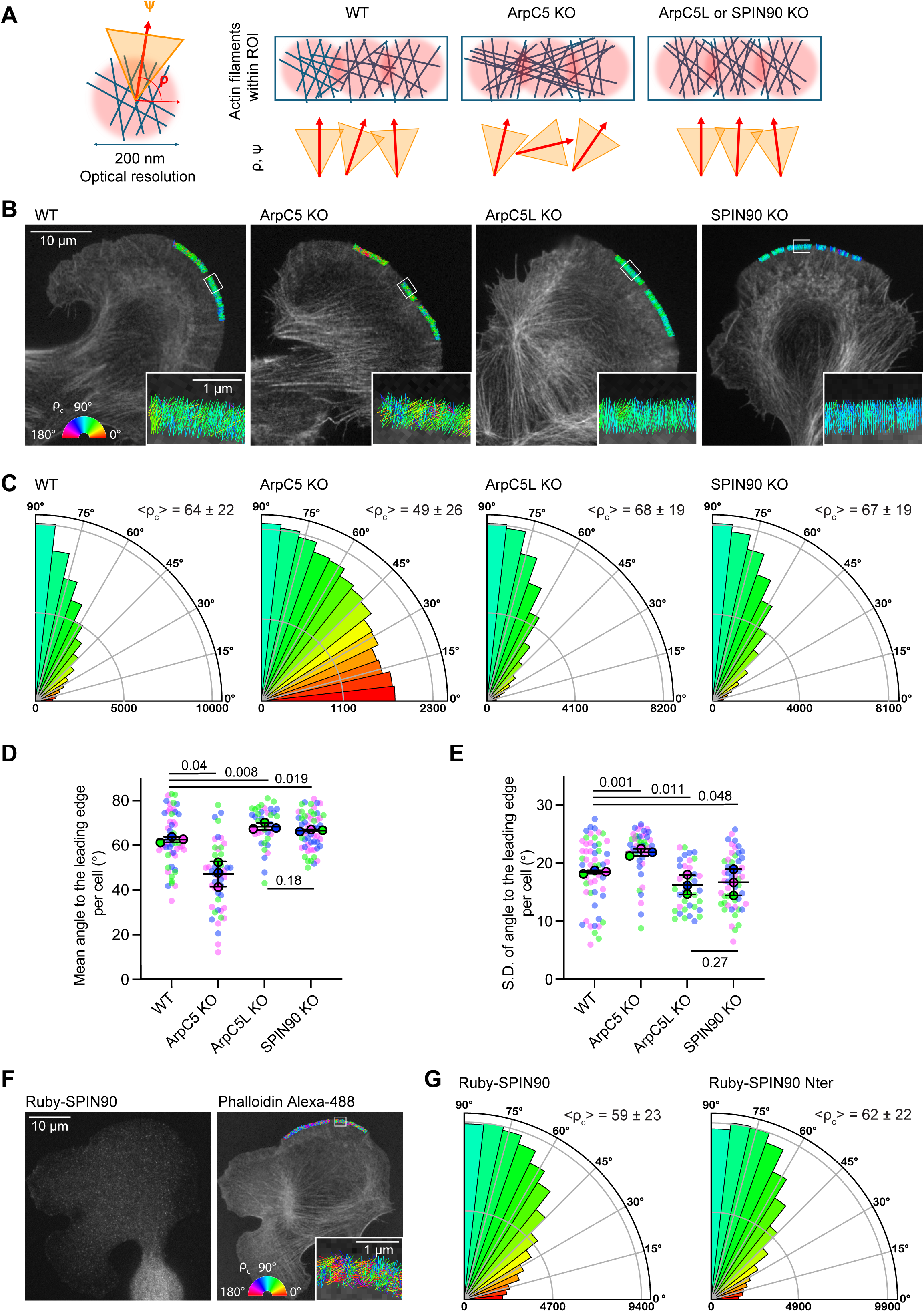
The organisation of actin filaments at the leading edge of B16 cells. (A) Schematic representation of ρ (rho) and ψ (psi) in the organization of the actin network. (B) Representative images of an Alexa-488 phalloidin stained B16 cell for each cell line as indicated. The coloured region in the lamellipodium is the result of polarimetry analysis within a 0.65 µm (10-pixel) region from the leading edge to determine the mean actin filament orientation in each pixel. The mean filament orientation per pixel is represented with a stick whose orientation corresponds to filament orientation (angle rho, ρ) and whose colour depicts filament orientation with respect to the leading edge (angle rho_c_, ρ_c_) within the 0–180° range according to the colour bar. The stick maps of a representative ROI (labelled with white rectangles) are zoomed in and shown. (C) The polar histogram shows the statistical distribution of actin filament angles (ρ_c_) per pixel within the 0–90° range relative to the leading edge of all the cells, with 38 to 55 cells analysed per condition across three independent repeats. The average < ρ_c_ > and standard deviation for each cell line are indicated. The colours correspond to the colour bar shown in panel A. (D) The statistical results of the average lamellipodial actin filament angle < ρ_c_ > relative to the leading edge within the 0–90° range in individual wildtype, ArpC5, ArpC5L or SPIN90 knockout cells. Each point represents the mean angle of actin filaments relative to the leading edge in a single cell from three independent experiments, the latter shown in different colours. Mean ± SD for each experimental condition is indicated. A two-tailed Welch’s t-test is used to estimate the p-value between the means under two conditions. (E) The statistical results of the standard deviation (SD) of actin filament angles relative to the leading edge within the 0–90° range in individual wildtype, ArpC5, ArpC5L, or SPIN90 knockout cells. Each point represents the SD of actin filaments relative to the leading edge in a single cell from three independent experiments, the latter shown in different colours. Mean ± SD for each experimental condition is indicated. A two-tailed Mann Whitney test is used to estimate the p-value between individual cells under two conditions. (F) Representative images of Alexa-488 phalloidin stained B16 cell (right) overexpressing Ruby-SPIN90 (left). The zoomed-in image shows the mean actin filament orientation per pixel within the ROI (white rectangle) as in panel A. (G) The polar histogram shows the statistical distribution of actin filament angles (ρ_c_) per pixel within the 0–90° range relative to the leading edge in all cells overexpressing either Ruby-SPIN90 (61 cells) or Ruby-SPIN90 Nter (59 cells), across three independent experimental repeats. The average < ρ_c_ > and standard deviation for each experimental condition are indicated. The colours correspond to the colour bar shown in panel C.

We measured high ψ values for lamellipodial actin (mean = 157°) in wildtype cells, as well as in all KO cell lines (Fig. S4A-C), with no detectable statistical difference among the different cell lines. This finding is in line with electron microscopy showing that lamellipodial actin filaments explore varying orientations within the scale of ∼200 nm (Fig. 6A, B), and with quantification of branch densities which were comparable in tomograms of WT, ArpC5 KO and ArpC5L KO cell lines (26).

Importantly, quantification of mean actin filament orientations showed distinct differences in our different cell lines (Fig. 6A-E and Fig. S4D). Whereas lamellipodial actin in WT B16 cells is enriched in populations more perpendicular to the leading edge (Fig. 6C, D), the distribution of filament orientations, both per pixel (Fig. 6C) and averaged per lamellipodium (Fig. 6D), is much broader in ArpC5 KO cells. This increased variability, reflected by the higher standard deviation (SD) of the respective distribution, indicates the coexistence of filament populations more parallel to the leading edge (Fig. 6E). Importantly, lamellipodial actin filaments in both ArpC5L KO and SPIN90 KO cells not only remained closer to perpendicular at the leading edge, but the respective distributions, both per pixel (Fig. 6C) and averaged per lamellipodium (Fig. 6D), were also narrower than in WT cells. This reflects the presence of a larger population of filaments approaching a right angle to the leading edge. Our observations are fully consistent with the quantification of individual filament angles from cryo-electron tomograms in ArpC5 isoform KO cells (26), and importantly further suggests that SPIN90 regulates lamellipodial actin architecture via ArpC5L.

Arp2/3-mediated branches follow a characteristic average 70-degree geometry whereas SPIN90–Arp2/3-nucleated linear actin filaments can in principle orient in all directions and also serve as mother filaments for new branches. Consequently, SPIN90-Arp2/3 is predicted to increase the diversity of filament orientations within a network. We therefore hypothesized that the absence of this linear filament population is responsible for the enrichment of filaments with angles perpendicular to the leading edge in both ArpC5L KO and SPIN90 KO cells. Moreover, this predicts a broader distribution of filament angles in cells overexpressing SPIN90. Consistent with this, we found that cells expressing Ruby-SPIN90 had a wider distribution of filament angles than those expressing Ruby-SPIN90 Nter which cannot interact with the Arp2/3 complex, the latter being more similar to wildtype cells (Fig. 6G, H and Fig. S4E, F).

### SPIN90 regulates lamellipodial protrusion

To examine the impact of the SPIN90 interaction with ArpC5L-containing Arp2/3 complexes on lamellipodial actin during protrusion, we generated B16 cells with endogenously tagged GFP-β-actin and lacking either the ArpC5 isoforms or SPIN90 respectively (Fig. 7A, S4A). As previously reported by Fässler et al. (2023), we observed that in B16 cells the absence of ArpC5 reduces lamellipodial width, while the loss of ArpC5L increases it (Fig. 7B). In contrast the SPIN90 KO cells have lamellipodia of similar width to those of wild-type cells (Fig. 7B). Quantification of the lamellipodial actin density (GFP-β-actin intensity), however, revealed that the actin density increased in SPIN90 KO cells compared to wild type controls. A similar increase in, lamellipodial actin density was also observed in the ArpC5L KO cells (Fig. 7B). In contrast, loss of ArpC5 resulted in reduced actin density. These results further confirmed that SPIN90 and ArpC5 isoforms have impact on the architecture of actin networks.

**Figure 7.**
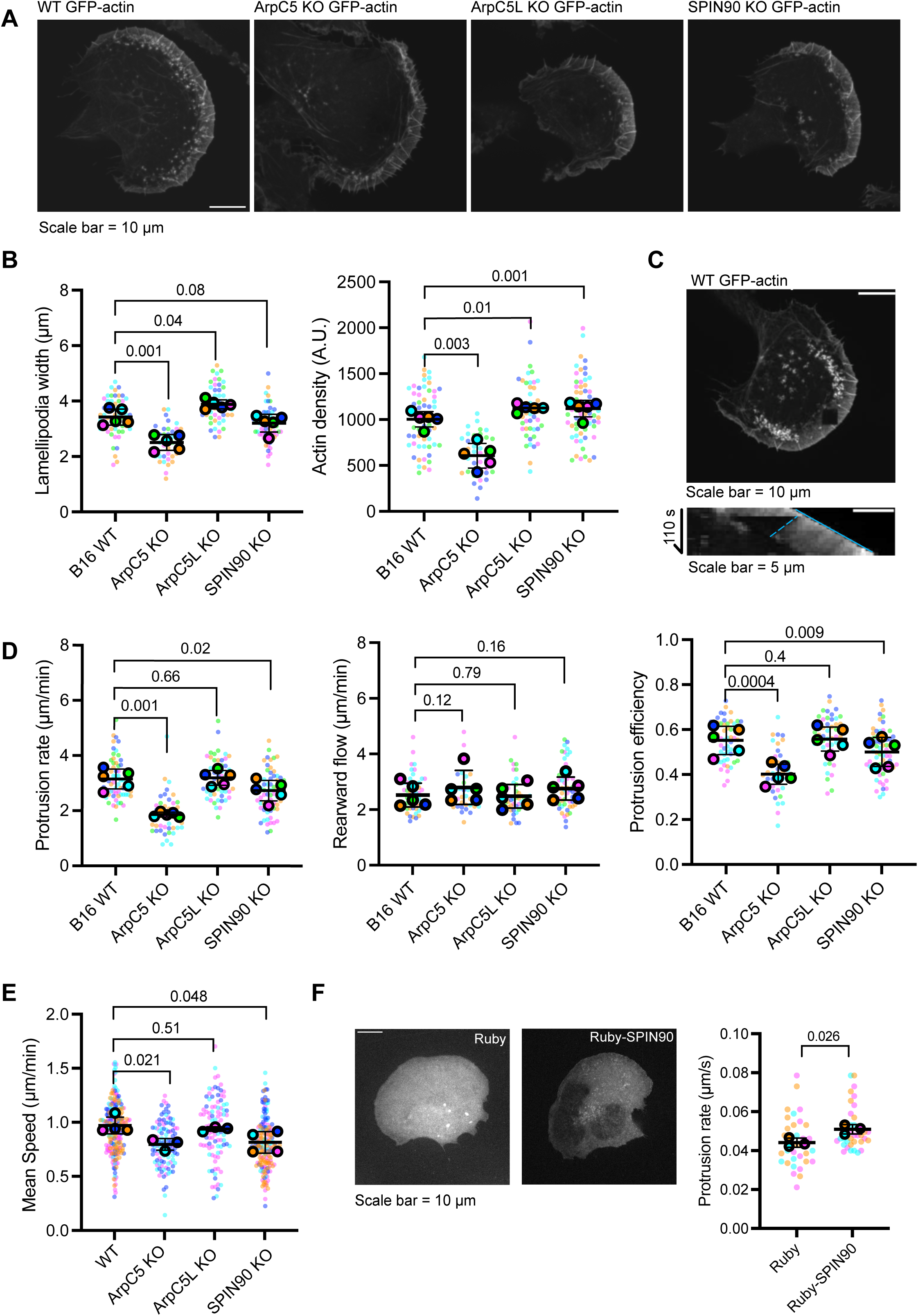
SPIN90 impacts the lamellipodia protrusion and cell motility. (A) Representative image stills of WT, ArpC5, ArpC5L and SPIN90 knockout B16 cells expressing endogenously tagged GFP β-actin (B) The quantification of lamellipodia width, lamellipodial actin density. Each point represents an individual cell, with different colours indicating independent experimental replicates. The mean of each experiment is shown as a circle, and error bars are standard deviation. A two-tailed paired t-test was used to estimate the p-value between the means under two conditions. (C) Representative image still (upper) from a FRAP assay on a B16 cell with endogenously GFP tagged β–actin. The kymograph (lower) shows the cell protrusion (indicated by straight line) and rearward flow (indicated by dashed line), made visible by the FRAP region. (D) The quantification of lamellipodial protrusion rate, rearward flow and protrusion efficiency. Each point represents an individual cell, with different colours indicating independent experimental replicates. The mean of each experiment is shown as a circle, and error bars are standard deviation. A two-tailed paired t-test was used to estimate the p-value between the means under two conditions. (E) Quantification of B16 cell random migration speed. Each point represents an individual cell, with different colours indicating 3 or 4 independent replicates. The mean of each experiment is shown as a circle, and error bars represent standard deviation. A two-tailed unpaired t-test was used to estimate the p-value between the means under two conditions. (F) Representative images (left panels) of the leading edge of B16 cells overexpressing either Ruby, or Ruby-SPIN90. The right panel shows the protrusion rate of these cells. Each point represents an individual cell, with different colours indicating independent experimental replicates. The mean of each experiment is shown as a circle, and error bars are standard deviation. A two-tailed paired t-test was used to estimate the p-value between the means under two conditions.

We performed FRAP analysis and quantified the protrusion rate and rearward flow of lamellipodial actin to examine the role of SPIN90 and ArpC5 isoforms on lamellipodial actin dynamics (Fig. 7C, D). Our results revealed that the loss of ArpC5 reduces the lamellipodial protrusion rate, while the absence of ArpC5L does not have a strong impact on protrusion. Nevertheless, we found that SPIN90 KO cells consistently exhibit a slower protrusion rate compared to controls (Fig. 7D). We did not observe any significant difference in rearward flow between the wild-type and the three KO cell lines. The sum of rearward flow and protrusion rate was considered the lamellipodial actin polymerization rate (Fig. S4B). When we calculated protrusion efficiency by dividing the protrusion rate by the actin polymerization rate, we found that ArpC5L KO cells maintain a protrusion efficiency comparable to wildtype cells, whereas both ArpC5 KO and SPIN90 KO cells have reduced protrusion efficiency (Fig. 7D). Consistent with this, in a random migration assay using unlabelled B16 cells, we observed that ArpC5 KO and SPIN90 KO cells migrated more slowly compared with wild-type and ArpC5L KO cells (Fig. 7E). We hypothesized that SPIN90–Arp2/3–nucleated linear actin filaments help support lamellipodial protrusion. To test this notion, we overexpressed Ruby-SPIN90 in B16 cells and found as predicted that Ruby-SPIN90 increased the lamellipodial protrusion rate (Fig. 7F).

## Discussion

Using in vitro biochemical approaches, we have now shown that the initiation of linear actin filaments by Arp2/3 complexes activated by SPIN90 is determined by the identity of their ArpC5 subunit. Only Arp2/3 complexes containing ArpC5L and not those with ArpC5, can be efficiently activated by SPIN90 (Fig. 1), due to their differing affinity for this nucleation promoting factor (Fig. 2). This makes SPIN90 the first Arp2/3 activator with a clear preference for a defined subset of Arp2/3 iso-complexes. Our findings further strengthen the hypothesis that Arp2/3 iso-complexes have distinct molecular and cellular properties determined by their subunit composition. Remarkably, while it is known that dimeric SPIN90 can bind two Arp2/3 complexes, our data fit a stoichiometric one-to-one binding model, leaving it unclear whether cooperative binding occurs between the complexes. We also found that SPIN90 is enriched at the leading edge of lamellipodia (Fig. 3B) where it enhances the recruitment of ArpC5L- but not ArpC5- containing Arp2/3 iso-complexes (Fig. 5). Importantly, we observed SPIN90 undergoes rearward flow towards the cell body (Fig. 4). Retrograde lamellipodial actin flow is known to result from myosin contraction and the resistive forces encountered during protrusion (45, 46). It is also established that after initiating a branch, the Arp2/3 complex remains at the branching point and becomes integrated into the branched actin network that undergoes rearward flow (47). The fact that SPIN90, but not its N-terminal domain which cannot bind Arp2/3, undergoes rearward flow indicates that SPIN90 and ArpC5L containing Arp2/3 integrate into lamellipodial actin together, presumably through crosslinking between SPIN90-Arp2/3-nucleated linear filaments and the rest of the lamellipodial actin.

Furthermore, using polarized fluorescence microscopy, we found that both SPIN90 and ArpC5 isoforms contribute to the architecture of lamellipodial actin. A likely scenario is that branches generated by Arp2/3, regardless of whether they contain ArpC5 or ArpC5L, are constrained by two factors: the orientation of the pre-existing mother filaments and the fixed 70° branching angle dictated by the structure of the Arp2/3 complex. In contrast, linear filaments generated by SPIN90–Arp2/3–ArpC5L are likely freer to orient before being crosslinked to actin filaments in the lamellipodia. In addition, these filaments may serve as mother filaments, providing additional sites for Arp2/3 mediated branching. The presence of SPIN90 and ArpC5L-containing complexes therefore increases the diversity of filament orientations at the leading edge. Hence, we observed less variable distribution of the ρ angle in both SPIN90 and ArpC5L KO cells (Fig. 6C-E).

Our observations of ArpC5 KO and ArpC5L KO lamellipodial actin networks using polarized fluorescence microscopy are consistent with previous reports based on high-resolution electron microscopy. Previous quantification of individual filaments showed that in ArpC5 KO cells, a larger number of filaments were oriented parallel to the leading edge, while in ArpC5L KO cells, more filaments were oriented perpendicular to the leading edge (26). Importantly, our results further demonstrate that SPIN90 works together with ArpC5L to regulate lamellipodial actin architecture. Although our approach does not provide information on the orientation of single actin filaments, it enables accurate measurements of the mean orientation and alignment of actin filaments in much larger areas and in many more cells than is feasible in electron microscopy-based approaches. The detailed organization of the actin networks in the absence of SPIN90, however, should still be examined by electron microscopy in future studies. Nevertheless, our approach offers the possibility of studying actin architecture through live imaging (43), thereby enabling simultaneous observation of cell behaviour and actin organization in the future.

SPIN90 has recently been shown to promote focal adhesion maturation in U2OS cells which form prominent stress fibres as well as increase their collective cell migration in a scratch assay (19). In our single cell migration assays, we consistently observed that depletion of SPIN90 led to a mild reduction in both the lamellipodial protrusion rate and individual cell migration speed (Fig. 7D, E). We believe that actin dynamics and architecture must be finely tuned to ensure efficient lamellipodial protrusion and cell migration. Accordingly, SPIN90 KO cells, which exhibit a narrower distribution of filament orientations (Fig. 6C-E) and narrower lamellipodia (Fig. 7B) display reduced protrusion efficiency. In contrast, despite having a lamellipodial architecture, including both filament orientation angles and their distribution similar to that of SPIN90 KO cells, ArpC5L KO cells maintain a protrusion rate and lamellipodial width comparable to those of wild-type cells. This may be attributed to the positive correlation between ArpC5- nucleated branches and Ena/VASP activity, which was previously reported to promote rapid actin filament elongation (26). Moreover, the low protrusion speed and motility observed in ArpC5 KO cells are likely the result of a combination of their negative correlation with Ena/VASP and an inefficient actin architecture, in which many filaments are likely oriented toward the leading edge at shallow angles. Here we illustrate the importance of actin architecture on the lamellipodia protrusion efficiency. Nevertheless, a systematic understanding of how actin architecture and dynamics impact protrusion efficiency, particularly under different external mechanical conditions remains to be established. Based on our observations and those of Pollard (19), SPIN90 appears to play a dual and seemingly opposing roles in cell migration: promoting cell protrusion while simultaneously restraining cell movement. The balance between these functions is likely to be finely regulated and may depend on the specific physiological context.

## Materials and Methods

### Protein purification and preparation

Recombinant human Arp2/3 iso-complexes with defined composition (Uniprot P61160, P61158, Q92747, O15143, O15144, O15145, P59998, Q9H9F9 and Q98PX5) were purified as previously reported (48). In this study, we focus on the Arp2/3 complex with Arp3 background because our previous study showed that both Arp3B and Arp3 containing Arp2/3 complex can generate linear filaments with SPIN90 (35). Skeletal muscle α-actin was purified from rabbit muscle following the published protocol (49). GST-tagged and untagged human SPIN90 C-terminus (267-715, UniProt Q9NZQ3-3) were purified as previously reported (50). Similarly, GST-tagged and untagged mouse SPIN90 full length (UniProt Q9ESJ4) were purified. Actin was fluorescently labelled on its lysine 328, with Alexa-488 succinimidyl ester (Life Technology) following the reported protocol (51). Actin was labelled with N(1-pyrene)-iodoacetamide (Thermo Fisher Scientific) as previously described (52).

### Pyrene actin polymerization assay

To probe actin polymerization, 100 nM SPIN90-Cter and 10 nM Arp2/3 iso-complexes were mixed with 2.5 μM Actin (5% pyrene labelled) and loaded immediately on a Safas Xenius fluorimeter. The negative control experiment was done without SPIN90-Cter. The experiments were performed at room temperature with a buffer containing 5 mM Tris-HCl pH 7.0, 0.1 mM CaCl2, 0.2 mM ATP, 50 mM KCl, 1 mM MgCl_2_, 0.2 mM EGTA and 1 mM DTT.

### TIRF assays

The coverslips were treated with Deep UV and passivated by mPEG silane overnight. The passivated coverslips were rinsed thoroughly with ethanol and water. Then they were dried and stuck to the glass slides with 3-mm-separated parafilm bands to create flow chambers. The experiments were performed at 25 °C. To begin with, 200 nM SPIN90-Cter, 20 nM Arp2/3 iso-complexes were mixed in a buffer containing 5 mM Tris-HCl pH 7.0, 50 mM KCl, 1 mM MgCl_2_, 0.2 mM EGTA, 0.2 mM ATP, 10 mM DTT, 1 mM DABCO, 0.1% BSA, and 0.3% Methylcellulose 4000 cP. The moment when 0.5 µM G-actin (15% Alexa-488 labelled) was added to the system was recorded as time 0. Negative control experiments were performed using the same system, either without SPIN90-Cter or with actin alone. For each condition, a random field of view was selected to quantify actin density. Actin density was quantified as the number of actin filaments divided by the surface area of the selected region.

### GST pull-down assays

The Glutathione Sepharose^TM^ 4B resin (GE Healthcare) was incubated with 5% BSA for 5 min and washed three times with GST pull-down buffer containing 50 mM Tris-HCl pH 7.0, 50 mM KCl, 1 mM MgCl_2_, 0.2 mM EGTA, 0.2 mM ATP, 1 mM DTT, and 5% glycerol. The beads were then incubated with 20 µM GST-SPIN90-Cter or 20 µM GST at 4 °C for 1h, followed by incubation with 100 nM Arp2/3 iso-complexes at 4 °C for 30 min. The beads were washed three times with 300 µL GST pull-down buffer. The proteins attached to the beads were eluted with 50 µL of 50 mM reduced glutathione. The sample was analyzed by SDS-PAGE followed by western blot using Anti-ArpC2 (sigma) to detect Arp2/3 complexes. Amersham imager AI600 or ImageQuant^TM^ 800 was used to develop the western blot results (Fig. S1D). To quantify the affinity between SPIN90 and Arp2/3 iso-complexes, we focused on Arp2/3-ArpC1B-ArpC5 and Arp2/3-ArpC1B-ArpC5L. Same amount of Arp2/3 complex (100 nM) was incubated with different concentration of GST bead attached SPIN90 Cter at RT for 1 hour. After centrifuging 500 g for 5 min, the same amount of supernatant was loaded on a SDS page gel. For a better sensibility, the stain free gel was visualized on a ChemiDoc imagining system, hence the subunits without Tryptophan, ArpC2 and ArpC4 are not visible. The fraction of bound Arp2/3 at a certain SPIN90 concentration equals 1-the intensity of ArpC1B divided by the intensity of ArpC1B when no SPIN90 added (Fig, 2A). The experiments were repeated independently three times and all the results were included in the plot. Alternatively, equal amounts of SPIN90-coated GST beads were incubated with varying concentrations of Arp2/3 iso-complexes at RT for 1 hour, The supernatant was removed and the beads were washed quickly with buffer. The attached component was eluted with 50 mM GSH. And the amount of eluted Arp2/3 was analyzed by western blot.

To quantify the dissociation rate of the Arp2/3 complex from SPIN90, equal amounts of SPIN90-coated GST beads were incubated with 2 µM Arp2/3 iso-complexes at RT for 1 hour. The supernatant containing unbound Arp2/3 complexes were removed after the centrifugation. The beads were then eluted at various time points with 50 mM GSH. The amount of eluted Arp2/3 was analyzed by western blot on a ChemiDoc imagining system. The blot is auto-exposed to improve resolution, as the exposure time for the sample containing ArpC5 is consistently much longer than that for ArpC5L. The experiments were repeated independently three times, and all the results were included in the final plot (Fig. 2B).

### Antibody, immunoblot analysis

The antibody against mouse SPIN90 (UniProt Q9ESJ4) was generated by immunizing rabbits with full-length purified recombinant mouse SPIN90 protein (WhiteAntibodies). Other antibodies are including: ARPC2/p34-Arc polyclonal (Millipore, 07-227), ARPC5 monoclonal (Synaptic Systems, 305011), ArpC5L (Proteintech, 22025-a-AP), β-actin (Abcam, ab179467), anti-mouse IgG-IRDye® 800CW (LI-COR Biosciences, 926-32210) and anti-rabbit IgG-IRDye® 680RD (LI-COR Biosciences, 926-68071) or anti-rabbit IgG (HRP) secondary antibody. Unless otherwise specified, western blot results were developed on LI-COR Odyssey CLx imaging system. FIJI with plugin GelAnalyzer was used to analyse the results.

### Cell culture and cell lines

WT M. Musculus B16-F1melanoma cells were gifts from Klemens Rottner lab (41). Unless otherwise specified in this project, cells were cultured in Gibco DMEM (ThermoFisher, 41966-029) with additional 10% FBS (CAPRICORN, FBS-LE-12A) and 1% Penicillin-Streptomycin (P/S, ThermoFisher, 15140122) at 37°C and 5% CO_2_. To detach the cells, cells were washed with PBS once and then treated with 0.25% Trypsin-EDTA (ThermoFisher, 25200056) at room temperature for 2-3 minutes.

ArpC5, ArpC5L and SPIN90 KO cell lines were generated by GRSPR-Cas9-mediated genome editing approaches. In each case, to generate a cut in the target gene, a pair of guide RNA were cloned intro pSpCas9(BB)-2A-Puro (Addgene, ID 48139) respectively and introduced into B16-F1 at the same time. The ArpC5 guide RNAs are ‘GACCGGGCTGGGCTCGCTAA’ and ‘GCGAGGTGGACTCGTGCCTA’. The guide RNAs against ArpC5L are ‘ATTCGTCGATATCCACGCGG’ and ‘CGGCAATATCCTTCCCGGCG’.

The SPIN90 guide RNAs are ‘GAGAGCCTAGAAGTCTGCAT’ and ‘GCTCCGTGGTGTCCATTGGA’. Cells were transfected using Fugene® HD Transfection Reagent. The next day, cells were selectively cultured by introducing 2.5 µg/mL puromycin into the medium. After 3 days, surviving cells were trypsinized and resuspended in PBS with additional 5% FBS. The single cells were sorted by the flow cytometry and seeded in DMEM with additional 20% FBS and 1% P/S in 96 well plates. Two weeks later, single colonies were expanded for characterizing, including Western blot and genomic sequencing of the target loci.

ArpC5 and ArpC5L genes were modified using ORANGE, a CRISPR-Cas9–mediated genome editing toolbox to fuse a NeonGreen tag with the target gene (53). pOrange NeonGreen-ArpC5 KI plasmid was modified based on pOrange GFP-ArpC5 KI (Addgene #131503). pOrange ArpC5L-NeonGreen KI plasmid was generated using the guide RNA: ‘CAGTGTGTTCCGGGCCATGG’. All the positive clones we tested (approximately 10 for each condition) were heterozygous with a higher fraction of endogenous ArpC5 isoforms remaining untagged (Fig. S2A, B). The same method was used to generate endogenously GFP-tagged β-actin. The plasmid pOrange GFP-Actb KI (Addgene #131479) was transfected into the WT, ArpC5 isoform KO, SPIN90 KO cells respectively. Single cells with green fluorescence were sorted by flow cytometry and seeded into 96-well plates containing DMEM supplemented with 20% FBS and 1% P/S. Expanded single colonies were examined by Western blot. All the positive clones we tested (approximately 10 for each condition) were heterozygous with a higher fraction of endogenous beta actin remaining untagged (Fig. S4A, right panel). B16 cells overexpressing GFP was generated using lentivirus infection (Trono group second generation packaging system, Addgene) and selected using puromycin resistance (1 µg/mL) as previously described (23).

### siRNA transfection

All-Star control (Qiagen, SI03650318) was used as the control. The following RNAi were purchased from Dharmacon (SMARTPools): siGENOME mouse ArpC5 (NM_026369), siGENOME mouse ArpC5L (NM_028809), siGENOME mouse NCKIPSD/SPIN90 (NM_030729). The final concentration of 20 nM was used. The cells were transfected with HiPerFect (Qiagen) using the fast forward method in 6-well plates. The experiments were performed 48h after the transfection. At the same time, the depletion of target gene was examined with Western Blot.

### Immunofluorescence on fixed cells

B16 cells were seeded on laminin coated coverslips as previously reported (26), and cultured in CO_2_ independent medium (Gibco^TM^) with additional 10% FBS and 2 mM L-Glutamine at 37°C. After 3h30 incubation, the cells were fixed with 4% PFA in PBS at 37°C. After fixation, cells were washed three times with warm PBS and permeabilized with 0.1% Triton X-100 in PBS for 1 min. Permeabilized cells were incubated with primary antibody for 40 minutes and secondary antibody for 40 minutes at room temperature (around 25°C). The samples were mounted with Mowiol (sigma). The coverslips were imaged on a Zeiss Axioplan2 microscope controlled with MetaMorph 7.8.13.0 software. A 63x/1.4 NA Plan-Achromat objective and a Photometrics Cool Snap HQ cooled charge-coupled device camera were used.

### Polarized fluorescence microscopy on fixed cells

B16 cells were seeded on laminin coated coverslips as previously reported (26), and cultured in CO_2_ independent medium (Gibco^TM^) with additional 10% FBS and 2 mM L-Glutamine at 37°C. After 3h30 incubation, the cells were fixed with 0.35% glutaraldehyde and 4% PFA in cytoskeleton buffer at 37°C. The samples were washed twice with PBS containing 1 mg/mL sodium borohydride and three times with PBS. The actin cytoskeleton was stained with 0.22 µM Phalloidin Alexa-488 at room temperature for 2 h in PBS containing 0.1% saponin and 10% BSA. The samples were mounted with Mowiol (sigma) and kept at 4 °C.

The images were acquired using a custom spinning disk (CSU-X1-M1; Yokogawa, Tokyo, Japan) polarization-resolved fluorescence microscope with a Nikon Plan Apo ×100/1.45 NA oil immersion objective lens (Ti2-E Nikon, Tokyo, Japan), 488- and 561-laser lines and an iXon Ultra 888 EMCCD camera (Andor, Belfast, UK), described in detail in Martins et al, 2025. The method is based on the rotation of the incident linear polarization in the plane of the sample, using an Electro Optics Modulator (EOM) made of a Pockels Cell (Pockels cell, No 28-NP; Quantum Technology, Lake Mary, FL), followed by a quarter wave plate (WPQ05M-488; Thorlabs, Newton, NJ). The microlens and pinhole array of the CSU spinning disks rotate synchronously, while the EMCCD and EOM are synchronized to ensure a fast stack recording for a given number of incident polarizations. To vary this incident polarization, voltage steps are sent to the EOM at the image acquisition rate. The linear polarization quality is assessed at the sample plane by measuring its residual degree of ellipticity - using a rotating quarter-wave plate followed by a polarizer and a photodetector-which is found to be less than a few percent. Polarization distortions are compensated by incorporating in the beam path a second dichroic mirror similar to the one present in the spinning disk (Di01-405/488/561/640; Semrock, Rochester, NY). Typically, polarized image stacks using 18 polarization angles with steps of 10° were recorded with an exposure time of 0.3-0.4 s per polarized image. A stack of images is recorded to extract the intensity modulation response of each pixel as a function of polarization rotation. The peak of this modulation yields the mean angle ρ of the fluorophores’ angular distribution within the focal volume of each pixel, while the modulation contrast corresponds to the angular width ψ of this distribution, which reflects the degree of molecular alignment within the fluorophore population. In practice, ρ and ψ are deduced from the measurement of the intensity modulation *I*(*α*), where *α* is the incident polarization angle. This is done by decomposing the dependence of *I*(*α*) as a modulation function *I*(*α*) = *a*_0_ + *a*_2_. cos 2*α* +*b*_2_. sin 2*α*. The coefficients (*a*_2_, *b*_2_) are determined by *a*_2_ = 2/*a*_0_. ∑*_k_ I*(*α_k_*). cos 2*α_k_* and *b*_2_ = 2/ *a*_0_. ∑*_k_ I*(*α_k_*). sin 2*α_k_*, where *a*_0_ = ∑*_k_ I*(*α_k_*) is the total intensity and *α_k_* are the angles used for the polarization-resolved measurements. The retrieval of (ρ, ψ) from the (*a*_2_, *b*_2_) coefficients is computed numerically, accounting for potential residual polarization distortions included in the model relating *I*(*α*) with the incident polarization *E*(*α*). In this model, *I*(*α*) follows the absorption polarization dependence such that *I*(*α*) ∝ ∫ ∫|*μ*(*θ*, *φ*). *E*(*α*)|^2^ sin *θ dθ dφ* where *μ*(*θ*, *φ*) = *μ*(cos *φ* sin *θ*, sin *φ* sin *θ*, cos *θ*) is the molecular absorption dipole oriented along the orientation (*θ*, *φ*) in the (*x*, *y*, *z*) sample frame and *E*(*α*) = (cos *α*, sin *α*, 0) is the incident polarization field.

Analysis and data representation, including color-coded stick representations of ρ angles with respect to the leading edge (ρ_c_) and of ψ angles per pixel, histograms and polar histograms were generated with the open-source PyPOLAR software (RRID:SCR_024681) (43). Pixels of the lamellipodia selected for analysis were identified using a combination of intensity thresholding and the “edge detection” tool in PyPOLAR to detect the leading edge and manual region selection, excluding microspikes. Only 10 pixels (0.65 µm) from the leading edge were included for analysis.

### Live cell imaging

The siRNA treated NeonGreen-ArpC5 and ArpC5L-NeonGreen cells were seeded on laminin-coated coverslips in Leibovitz’s L-15 medium supplemented with 10% FBS and 2 mM L-glutamine at 37 °C. After 3-hour incubation, the cells were imagined on a Zeiss Axio Observer spinning disc with a Plan Achromat 100x/1.46 NA oil lens, an Evolve 512 camera, and a Yokagawa CSUX spinning disk. The microscope was controlled by the SlideBook software (3i Intelligent imaging Innovations). For each condition, more than 8 cells with lamellipodia were chosen randomly. The same exposure time, 200 ms, was applied in each acquisition. The intensity of ArpC5 isoforms signal at the leading edge was quantified as the measured intensity at the leading edge minus the intensity of the background. The normalised intensity of ArpC5 isoforms signal was the intensity of ArpC5 isoforms divided by the expression level of NeonGreen tagged ArpC5 quantified by the western blots.

Similarly, WT and KO cells expressing endogenous GFP-β-actin were seeded and imagined on the confocal spinning disc with a Plan Achromat 100x/1.46 NA oil lens. Life images were acquired at 5 second intervals. A region of interest on the lamellipodia was photobleached using a focused 3-ms pulse of a 405 nm laser. Images were analyzed using FIJI software with Multi Kymograph. For each individual cell, the protrusion rate and the rearward flow were measured respectively (Fig. 7C, D). The polymerization rate is the sum of protrusion rate and rearward flow (Fig. S4B). The protrusion efficiency was quantified as the protrusion rate divided by the polymerization rate (Fig. 7D).

Ruby-tagged SPIN90 was transiently introduced into cells using the FuGENE kit 48 hours before the experiment. The ArpC5L-NeonGreen or GFP overexpressing Ruby-tagged SPIN90 cells were seeded on laminin-coated coverslips in Leibovitz’s L-15 medium supplemented with 10% FBS and 2 mM L-glutamine at 37 °C. After 3 hours, the cells were imagined using instant Structured Illumination Microscopy (VT-iSIM). VT-iSIM was conducted on an Olymus iX83 Microscope with Olympus 150x/1.45 NA X-Line apochromatic objective lens, dual Photometrics BSI-Express sCMOS cameras, and CoolLED pE-300 light source (Visitech), controlled using Micro-Manager. Images were acquired at 1 second intervals with 200 ms exposure time. Images were analyzed using FIJI software. Normalized fluorescent signal intensity at the leading edge was quantified by dividing the intensity at the leading edge (solid line, Fig. 3A) by the intensity measured 2 µm behind the leading edge (dashed line, Fig. 3A).

To quantify the random motility, WT and CRISPR KO cells were imaged and analysed using the Livecyte system (PhaseFocus) every 10 minutes for 16 hours at 10x. The imaging chamber was maintained at 37 °C with 5% CO_2_. Livecyte automatically process the data and quantified the cell motility.

### Protein structure analysis

Pymol is used to visualise and analyse the structure and electron density maps of SPIN90 bound Arp2/3 (54).

### Statistical analysis

Statistical analyses were performed with Prism 10 (Version 10.1.1). Sample numbers, mean and P value can be found in the figures or figure legends. Unless otherwise specified, each experiment have been independently repeated three times. P-values smaller than 0.05 are considered statistically significant. Moreover, Cohen’s d was also used to assess the effect size; a Cohen’s d greater than 0.8 is considered a large effect.

Cells receiving the treatment (e.g., siRNA) on the same day were paired. A paired two-tailed Student’s t-test was then used to compare the means between the two groups. Also, a paired two-tailed Student’s *t*-test was used to compare the signal intensity of GFP and Ruby SPIN90 at the leading edge. Because the data distribution in the polarized microscopy assay differs substantially, we used an unpaired two-tailed *t*-test with Welch’s correction or Mann Whitney test to compare the two groups. A paired two-tailed *t*-test with was used to compare lamellipodia protrusion between wild type and KO cell lines.

## Supplementary

### Sequence results

<u>The grey colored fragment is present in WT cell lines but missing in KO cells. B16-F1 ArpC5 knockout lines</u>

Reference: MmC5 WT CCGGGCTGGGCTCGCTAAAGGAGAGGCACCGCGGAGGGGCAGGCCAGCGTCGCGTCGCGATCCGGGATGTCGAAGAACACGGTGTCGTCGGCCCGCTTCCGGAAGGTGGACGTGGACGAATATGACGAGAACAAGTTCGTGGACGAGGAGGACGGCGGCGATGGCCAGGCCGGGCCCGACGAGGGCGAGGTGGACTCGTGCCTACGGCAATATC

MmC5 KO 2.1 CCGGGCTGGGCTCGCGCGGCAATATC

B16-F1 ArpC5L Knockout lines

Reference: MmC5L WT

CGCTTCCGCCGCGTGGATATCGACGAATTTGACGAGAACAAATTCGTAGACGAGCACGAAGAGGCAGCGGCGGCGGCGGGCGAGCCAGGCCCCGACCCCTGCGAGGTAGACGGGCTCCTGCGGCAATATCCTTCCCGGCGCGGCGGCCCGGCT

MmC5L KO 2.2 CGCTTCCGCCGCGTGGCGCGGCGGCCCGGCT

B16-F1 SPIN90 Knockout lines

Reference: SPIN90 WT

GAGGATGGCCTTCCAATGGACACCACGGAGCAGCTGCCAGACCTCTGCATGAACCTGCTTCTGGCTCTCAACCTGCACCTGACAGGTGGGGTGGGGCCATCTGGAAGGCAGCTGAGCAGTGGGCTTTTCGGGCTTCGCGGGCTTCTGTGCCTACACTCTTACCTGTCTCCATCCACGTCTCTCCGCAGCTCCTGAGCAGAATGTCATCATGGCTGCCTTGAGCAGACACACCAATGTGAAGATCTTCTCTGAGAAGCTGCTTCTGCTTCTGAACAGAGGGGGTGAGAGCCTAGAAGTCTGCATGGGTGCCGCTTGCGT

SPIN90 KO 2.10 GAGGATGGCCTTCCCATGGGTGCCGCTTGCGT

## Acknowledgements

This project has received funding from the European Research Council (ERC) under the European Union’s Horizon 2020 research and innovation programme (grant agreement No 810207 to M.W.). L.C. was supported by the European Union’s Horizon 2020 Marie Sklodowka-Curie individual fellowship program (H2020-MSCA-IF-101028239 – MolecularArp) and the French National Research Agency (ANR) grant ArpRegulation (ANR-25-CE13-0450). A.J. was supported by the Fondation Bettencourt-Schueller (Impulscience program). G. R.-L. was supported by the Fondation pour la Recherche Medicale (grant EQU202203014630). MW is supported by the Francis Crick Institute, which receives its core funding from Cancer Research UK (CC2096), the UK Medical Research Council (CC2096), and the Wellcome Trust (CC2096). M.V. and S.B. are supported by funding from ANR grant Equipex+ IDEC (France 2030 investment plan ANR-21-ESRE-0002) and the France-BioImaging infrastructure (ANR-10-INBS-04). We thank Tianyang Liu (Birkbeck College, UK) for comparing the published electron density map with different structural models. We thank Jean-Jacques Honorine (Institut Fresnel, France) for technical help with the calibration of the polarized fluorescence microscope. We acknowledge the Photonics facility of the Institut Fresnel. We thank Jeremy Carlton (King’s College London, UK) and Klemens Rottner (Helmholtz Centre for Infection Research, Germany) for feedback on the manuscript. For the purpose of Open Access, the authors have applied a CC BY public copyright licence to any Author Accepted Manuscript version arising from this submission.

## Supplementary Figure legends

**SF 1. Binding between the Arp2/3 complex and SPIN90.**

(A) Sequence alignment between ArpC5 and ArpC5L subunit. The interface binding SPIN90 is highlighted with colours: residues that are identical in both isoforms are shown in yellow, while distinct residues are shown in cyan.

(B) Surface representation of the Arp2/3 complex from the SPIN90-Arp2/3 complex structure (PDB: 6DEC). Residues in Arp2/3 complex subunits contacting the two SPIN90 molecules are shown in darker colours.

(C) The 2Fo-Fc electron density map of Arp2/3 crystal structure in complex with SPIN90 (purple) is shown at a contour level of 1.5σ (blue mesh). ArpC5L (dark yellow) is superimposed with ArpC5 (light yellow) subunit determined in the structure. The remaining subunits of Arp2/3 complex are coloured in grey.

(D) Immunoblotting using an ArpC2 antibody demonstrates that, when equal amounts of Arp2/3 are incubated with SPIN90, more ArpC5L-containing Arp2/3 complexes are pulled down with GST-SPIN90 than ArpC5-containing complexes.

(E) Immunoblots using an ArpC2 antibody show the amount of Arp2/3 complex containing ArpC1B pulled down with GST-SPIN90. Both blots were run on the same gel and developed on the same membrane to allow a direct comparison of signal intensity. The graph shows the quantification of average normalized ArpC2 intensity from three independent replicates. The data was fit with one site-specific binding equation.

**SF 2. ArpC5L and SPIN90 can localise and move together in the cell.**

Corresponding to Fig 4B. Another representative image of B16 cells with endogenously labelled NeonGreen-ArpC5L and overexpressing Ruby-SPIN90 at a certain time, defined as time 0. Right panel shows zoomed-in images of the several ROI highlighted in the orange box in the left panel. Yellow arrows indicate the exact position where ArpC5L and SPIN90 signals first observed overlap, while orange arrows show the ArpC5L and SPIN90 signals moving together directionally over time.

**SF 3. The western blot of siRNA treated cells.**

(A) The western blot of siRNA treated ArpC5-NeonGreen B16 cells.

(B) The western blot of siRNA treated NeonGreen-ArpC5L B16 cells.

(C) Representative immunofluorescence images of siRNA-treated B16 cells. The average ratio of ArpC5 to ArpC2 signal in AllStar control is defined as one. For each repeat, the ratio of ArpC5 to ArpC2 signal is normalized to the AllStar control. Quantitative results are shown in the right panel. Each dot represents an individual cell, and the circles indicate the mean of each repeat. The Wilcoxon test was used to examine statistical significance.

**SF 4. The organisation of lamellipodial actin.**

(A) Corresponding to Fig 6A, B. Representative images of an Alexa-488 phalloidin stained B16 cell for each cell line as indicated. The mean filament orientation per pixel is represented with a stick whose orientation corresponds to filament orientation (angle rho, ρ) and whose colour depicts filament alignment (angle psi, ψ) according to the colour bar. The stick maps of a representative ROI (labelled with white rectangles) are zoomed in and shown.

(B) The histogram shows the statistical distribution of filament alignment angles ψ per pixel of all the individual cells, with 38 to 55 cells analysed per condition across three independent repeats. The average <ψ> and SD for each cell line are indicated. The colours correspond to the colour bar shown in panel A.

(C) Statistical analysis of the average <ψ> in each individual wildtype, ArpC5, and ArpC5L or SPIN90 knockout cells. Mean ± SD for each cell line is indicated. A two-tailed Welch’s t-test is used to estimate the p-value between individual cells under two conditions.

(D) Corresponding to Fig 6C. The polar histogram shows the statistical distribution of actin filament angles (ρ_c_) per pixel within the 0–180° range relative to the leading edge in all cells. The average ρ_c_ per pixel and SD for each cell line are indicated. The colours correspond to the color bar shown in panel Fig 6A.

(E) Corresponding to Fig. 6G. The polar histogram shows the statistical distribution of actin filament angles (ρ_c_) per pixel within the 0–180° range relative to the leading edge in all cells, overexpressing either Ruby-SPIN90 or Ruby-SPIN90 Nter. The average < ρ_c_ > and standard deviation for each experimental condition are indicated.

(F) The statistical results of the average lamellipodial actin filament angle relative to the leading edge within the 0–90° range in individual wildtype cells (also shown in Fig. 6G) or cells overexpressing either Ruby-SPIN90 or Ruby-SPIN90 Nter. Each point represents the mean angle of actin filaments relative to the leading edge in a single cell from three independent experiments, the latter shown in different colours. Mean ± SD for each experimental condition is indicated. A two-tailed Welch’s t-test is used to estimate the p-value between the means under two conditions.

**SF 5. Analysis of KO cell lines.**

(A) The left panel shows the absence of targeted gene in different Crispr engineered cell lines, while the right panel show the endogenous labelling of GFP-β-actin in these cell lines.

(B) The graphs show quantification of the polymerization rate which is calculated by summing the rearward flow and protrusion rates (Fig. 7D). Each point represents an individual cell, with different colours indicating independent experimental replicates. The mean of each experiment is shown as a circle, and error bars indicated the standard deviation. A two-tailed paired t-test was used to calculate the p-value between the means under two conditions.

## Movies

**Movie 1**. Movie showing live imaging of B16 cells overexpressing GFP (a volume marker) and Ruby-SPIN90, corresponding to Fig. 3B (middle panel). The right panel shows inverted-intensity live images of each channel.

**Movie 2**. Movie showing live imaging of B16 cells overexpressing GFP (a volume marker) and Ruby-SPIN90 Nter, corresponding to Fig. 3B (lower panel). The right panel shows inverted-intensity live images of each channel.

**Movie 3**. Movie showing live imaging of NeonGreen-ArpC5L B16 cells overexpressing Ruby-SPIN90, corresponding to Fig. 4A (upper panel). The middle and the right panel show inverted-intensity live images of each channel, where several examples of moving NeonGreen-ArpC5L and Ruby-SPIN90 were shown by cyan circles respectively. One example of colocalised NeonGreen-ArpC5L and Ruby-SPIN90 were highlighted by yellow circle in the left panel.

**Movie 4**. Movie showing live imaging of NeonGreen-ArpC5L B16 cells overexpressing Ruby-SPIN90, corresponding to Fig. 4A (lower panel). The middle and the right panel show inverted-intensity live images of each channel. Several moving NeonGreen-ArpC5L were highlighted by cyan circles. No Ruby-SPIN90 was observed moving directionally.

