## supplemental figures 1 to 5 for "SPIN90 modulates the architecture of lamellipodial actin in an ARPC5L dependent fashion"

**A**

```
Hu_ArpC5         1 MSKNTVSSARFRKVDVDEYDENKFVDEEDGGDGQA---GPDEGEVDSCLRQGNMTAALQA
Hu_ArpC5L        1 MARNTLSS-RFRRVDIDEFDENKFVDEQEEAAAAAAEPGPDPSEVDGLLRQGDMLRAFHA
                   *  ** ** *** ** ** ********       *   ***  ***  **** *  *  *

Hu_ArpC5        58 ALKNPPINTKSQAVKDRAGSIVLKVLISFKANDIEKAVQSLDKNGVDLLMKYIYKGFESP
Hu_ArpC5L       60 ALRNSPVNTKNQAVKERAQGVVLKVLTNFKSSEIEQAVQSLDRNGVDLLMKYIYKGFEKP
                   ** * * *** **** **   *****  **   ** ****** *************** *

Hu_ArpC5       118 SDNSSAMLLQWHEKALAAGGVGSIVRVLTARKTV
Hu_ArpC5L      120 TENSSAVLLQWHEKALAVGGLGSIIRVLTARKTV
                     **** ********** ** *** *********
```

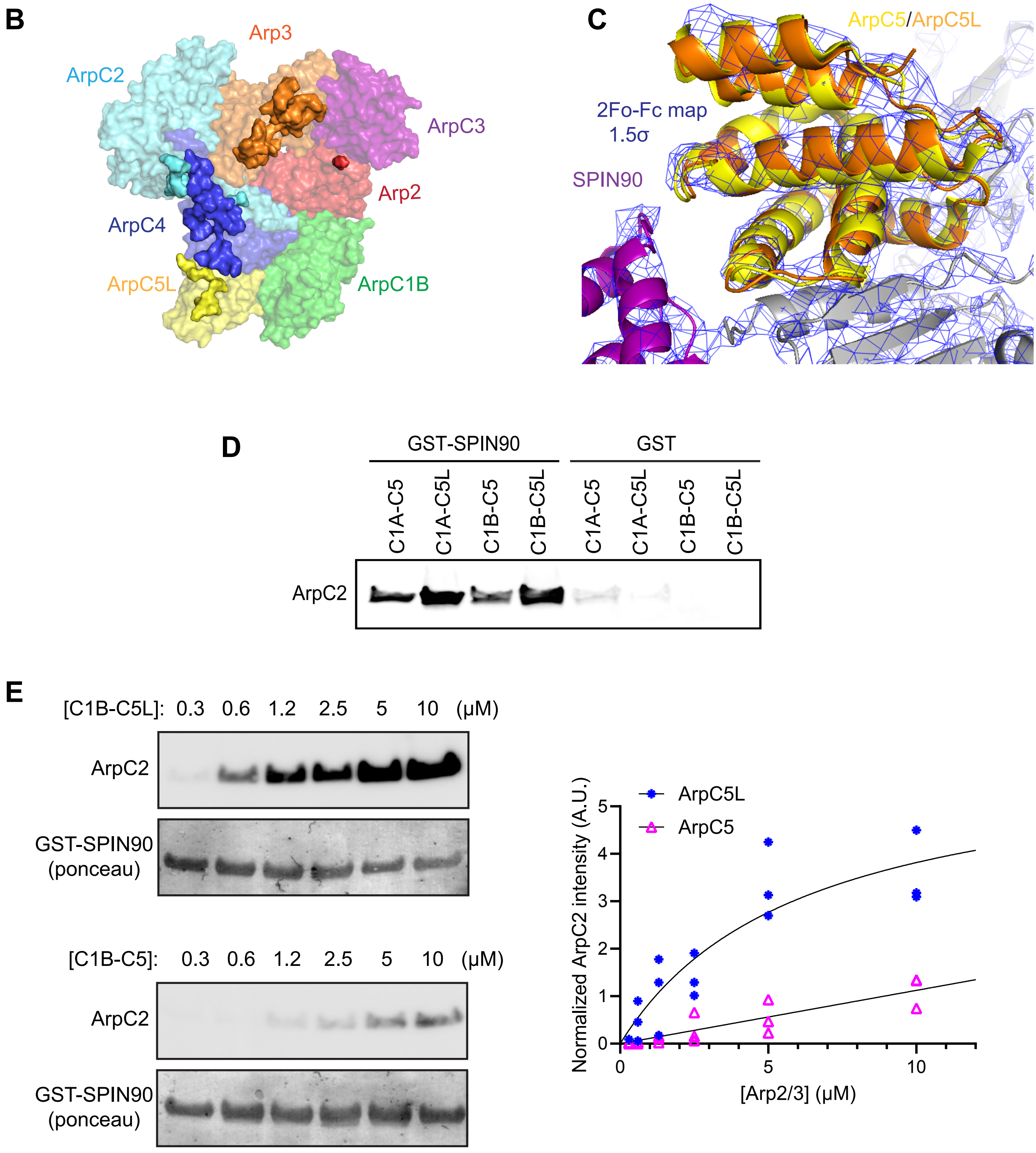


Supplementary Figure 1

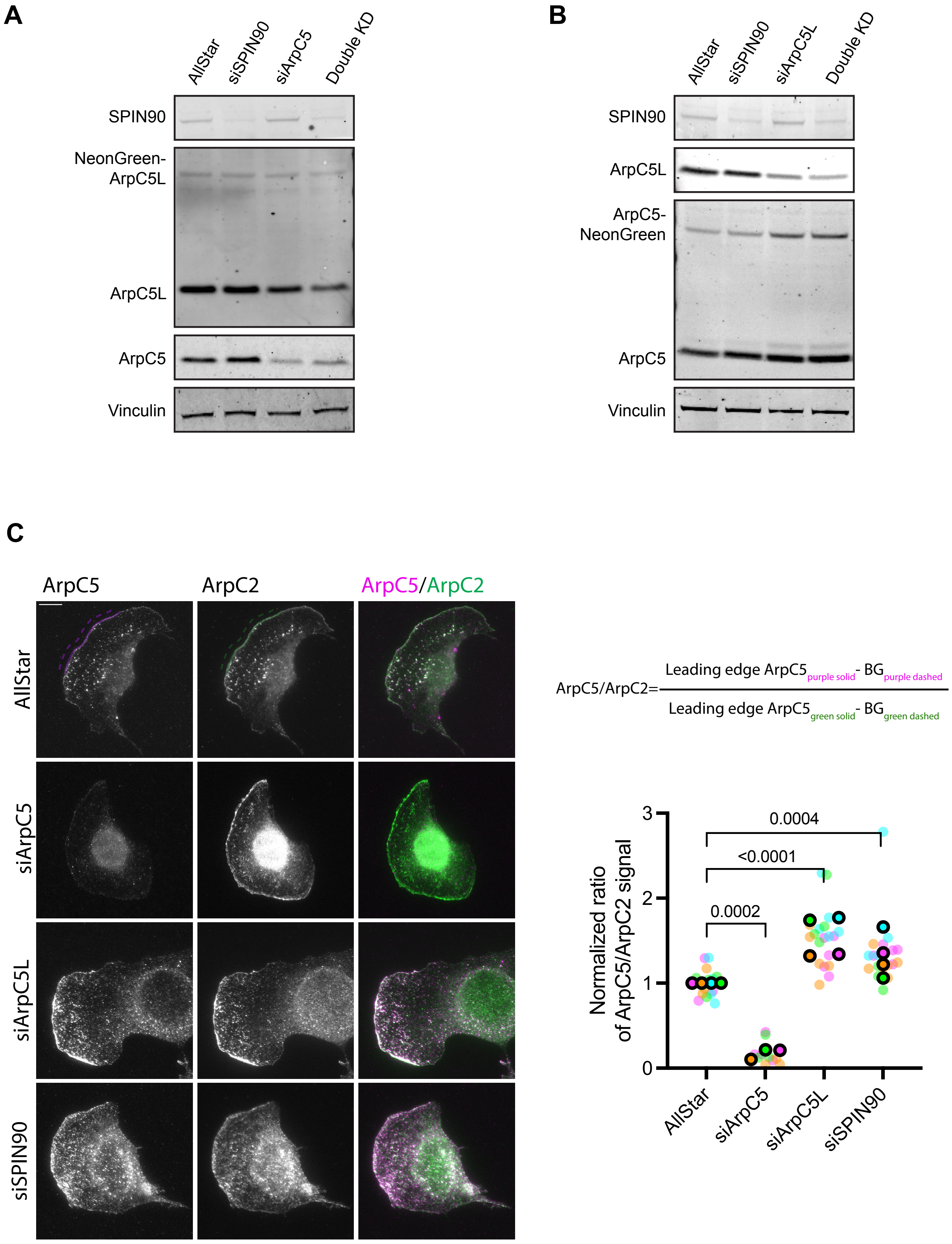


Supplementary Figure 2

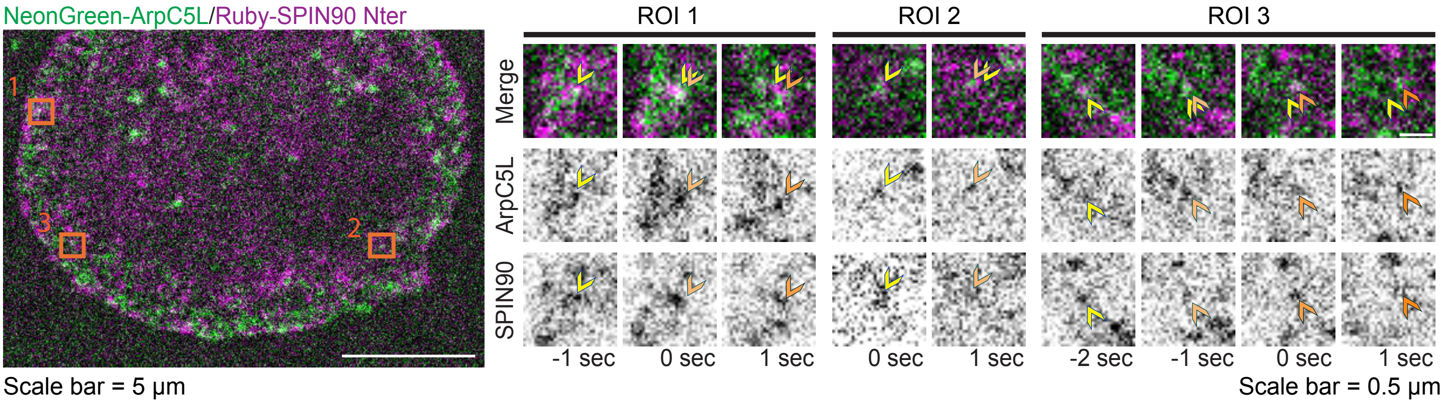


Supplementary Figure 3

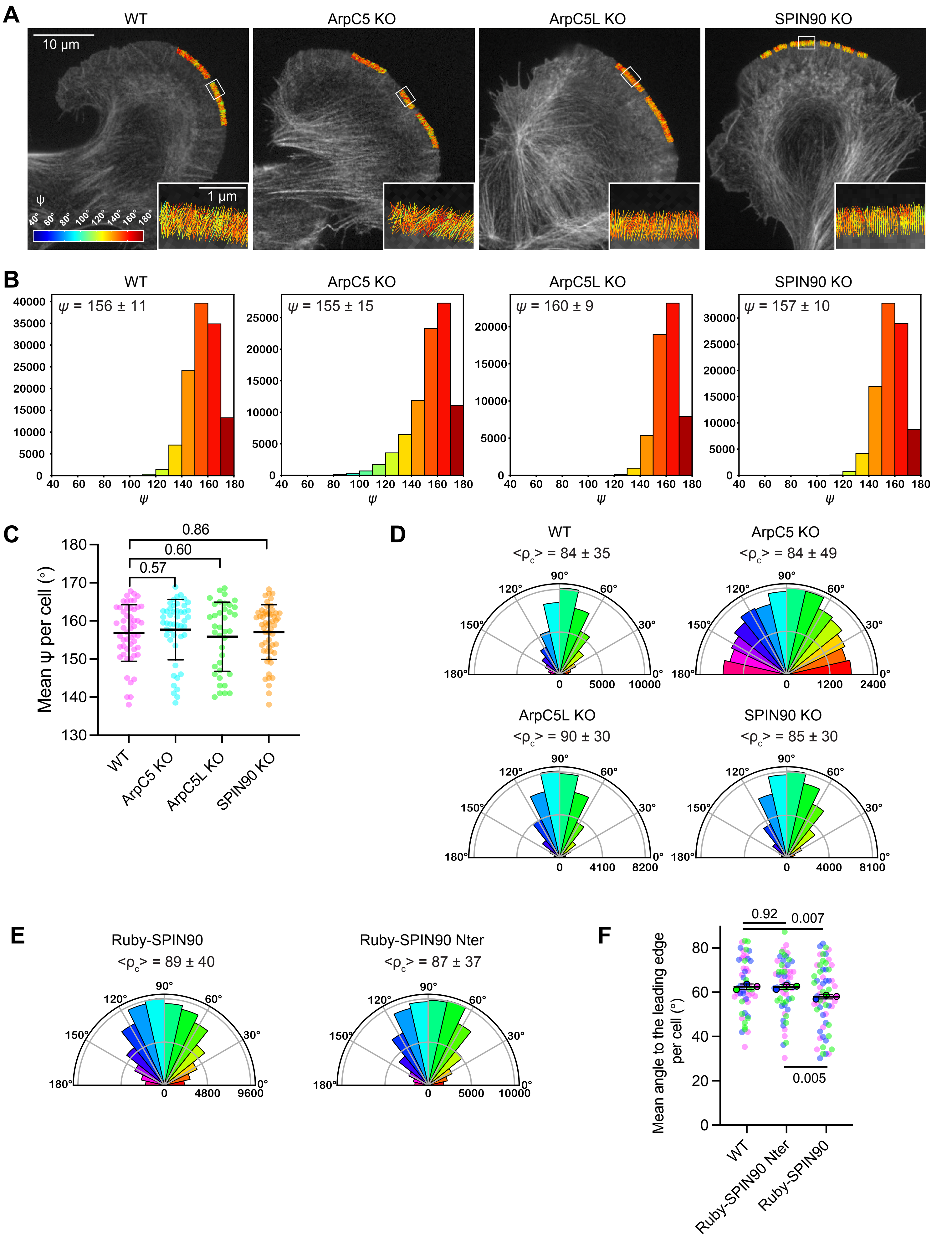
A
WT
ArpC5 KO
ArpC5L KO
SPIN90 KO
10 µm
1 µm
ψ
B
ψ = 156 ± 11
ψ = 155 ± 15
ψ = 160 ± 9
ψ = 157 ± 10
C
Mean ψ per cell (°)
0.86
0.60
0.57
D
<ρc> = 84 ± 35
<ρc> = 84 ± 49
<ρc> = 90 ± 30
<ρc> = 85 ± 30
E
Ruby-SPIN90
Ruby-SPIN90 Nter
<ρc> = 89 ± 40
<ρc> = 87 ± 37
F
Mean angle to the leading edge per cell (°)
0.92
0.007
0.005

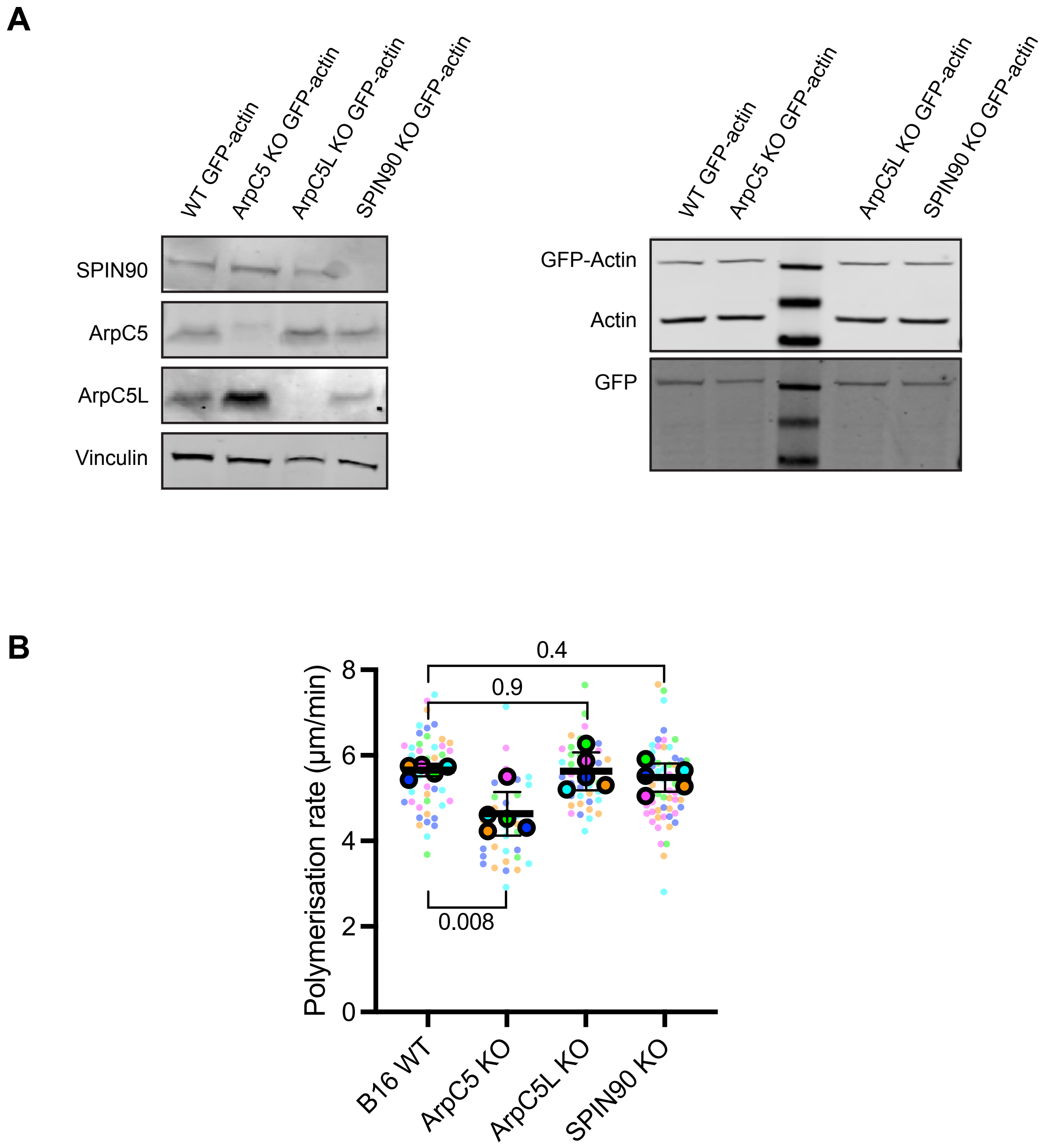


Supplementary Figure 5
